# Imaging Intrinsic Stochastic Magnetic Fluctuations in Living Cells

**DOI:** 10.64898/2026.03.23.713642

**Authors:** Weiming Lin, Tao Ding, Changming Bao, Yan Miao, Jiaxuan Zhou, Zhongxia Wei, Sisi Jia, Chunhai Fan, Le Liang

**Author notes:** These authors contributed equally to this work (Weiming Lin, Tao Ding).

## Abstract

**Significance:** This work establishes a probabilistic magnetometry framework for detecting weak stochastic magnetic fluctuations at the nanoscale, which provides the first quantitative access to intrinsic magnetic activity in living cells.

Weak stochastic magnetic fluctuations at nanoscale are difficult to quantify. In living cells, ionic transport and molecular currents generate electromagnetic activity whose magnetic component is likewise nanoscale, weak, stochastic, and rapidly varying and has therefore remained experimentally inaccessible. Here we introduce Bio-Spin Probabilistic Inference (BISPIN), a digital statistical framework that can quantify weak, stochastic magnetic fluctuations at the nanoscale. Using threshold-resolved signals from enhanced nitrogen-vacancy quantum sensors, BISPIN converts unstable analog magnetic readouts into statistically convergent digital observables and infers fluctuation strength through probabilistic modeling, enabling robust quantification under random sensor orientations and biological heterogeneity within the experimental bandwidth. Applied to living cells, this approach distinguishes live from fixed cells, resolves agonist-induced activation, and maps subcellular variations in magnetic fluctuation strength. By providing the first quantitative access to intrinsic stochastic magnetic fluctuations in living cells, this work establishes a probabilistic magnetometry framework for cellular electrodynamics and opens a new magnetic dimension of cellular phenotyping for bio-spin omics.

## 1. Introduction

Quantitative measurement of weak stochastic magnetic fluctuations remains a longstanding challenge at the nanoscale. Conventional magnetic sensing approaches are highly effective for static, coherent, or spectrally structured fields but become unreliable when signals are extremely weak, rapidly varying, and randomly oriented (*1, 2*). Within experimentally accessible bandwidths, such stochastic nanoscale magnetic signals are more appropriately characterized by their fluctuation strength than by a stable mean field. However, analog amplitude and spectral readout schemes are poorly suited to this regime, particularly when signals approach the noise floor and when environmental heterogeneity further complicates quantitative interpretation (*3, 4*).

Nitrogen-vacancy (NV) centers in nanodiamond (ND) provide a promising platform for magnetic sensing under ambient conditions at the nanoscale (*5–11*). However, conventional NV-based magnetometry detects only the magnetic-field projection along a given NV orientation (*12*). Weak stochastic signals with time-varying directionality do not necessarily yield stable resonance shifts that can be straightforwardly fit or tracked, and random sensor orientations further amplify this problem (*13, 14*). This creates a fundamental mismatch with analog magnetometry, which is optimized for deterministic magnetic observables. Moreover, translating NV magnetometry into a robust method for probing weak stochastic magnetic activity requires overcoming multiple limitations simultaneously, including insufficient photon budget and readout contrast in the weak-signal regime, instability of analog spectral fitting under stochastic perturbations, and uncertainty arising from random sensor orientation, optical drift, and environmental heterogeneity.

Living cells represent one of the most important and challenging systems in which this problem arises. Cells are intrinsically nonequilibrium electrodynamic systems (*15, 16*). Ionic transport across membranes, molecular charge displacement, and intracellular current flows continuously generate nanoscale electromagnetic fluctuations (*17, 18*). Electrical aspects of these processes have been extensively studied through patch-clamp recording, voltage imaging, and ion-sensitive fluorescence probes (*19–21*). In contrast, the magnetic component of cellular electrodynamics remains largely unexplored. In the experimental regime considered here, intrinsic cellular magnetic activity is more appropriately treated as stochastic magnetic fluctuations than as stable directional fields. As a result, direct experimental access to intrinsic cellular magnetic fluctuations has remained difficult.

Here we introduce Bio-Spin Probabilistic Inference (BISPIN), a digital statistical framework that enables quantitative access to intrinsic stochastic magnetic fluctuations at the nanoscale. Rather than extracting information from analog amplitude or spectral shifts, BISPIN converts stochastic magnetic signals into digital threshold-resolved events recorded by plasmon-enhanced NV quantum sensors. By modeling the probabilistic statistics of these threshold events, the method infers magnetic fluctuation strength within the experimental measurement bandwidth. This event-based strategy is intrinsically robust to random sensor orientation, biological heterogeneity, and slow instrumental drift, thereby transforming a weak and stochastic magnetic signal into a quantifiable observable.

To support spatially resolved probabilistic measurements, we implement BISPIN using a high-density plasmonic NDs sensing array that enables wide-field acquisition of threshold-resolved magnetic events across the imaging plane. Plasmonic enhancement improves photon collection efficiency and ODMR contrast while maintaining nanoscale proximity to living cells. Controlled random-field calibration establishes the quantitative relationship between event statistics and magnetic fluctuation strength, and independent controls confirm magnetic specificity of the inferred signals.

Using BISPIN, we quantitatively distinguish live from fixed cells, resolve agonist-induced electrodynamic activation, and map subcellular variations in stochastic magnetic fluctuation strength. Across multiple cell types, the resulting fluctuation maps yield reproducible cell-state fingerprints, revealing structured magnetic signatures associated with cellular activity. By providing quantitative access to intrinsic stochastic magnetic fluctuations in living cells, this work establishes a general measurement framework for cellular electrodynamics. More broadly, it introduces a new quantitative dimension for studying living matter, extending magnetic sensing from static field detection to stochastic electrodynamic processes at the subcellular scale.

## 2. Results and discussion

### 2.1 Probabilistic inference for stochastic nanoscale magnetometry

Weak stochastic magnetic perturbations at the nanoscale do not yield stable analog observables that can be straightforwardly quantified by deterministic field amplitudes. In NV-based sensing, only the magnetic-field projection along a given NV axis contributes to the measured resonance response. When magnetic perturbations vary in both magnitude and direction over time, the resulting instantaneous projections become strongly mixed, making direct analog estimation of field amplitude intrinsically unstable within the experimental bandwidth (Figure 1A, top).

**Figure 1.**
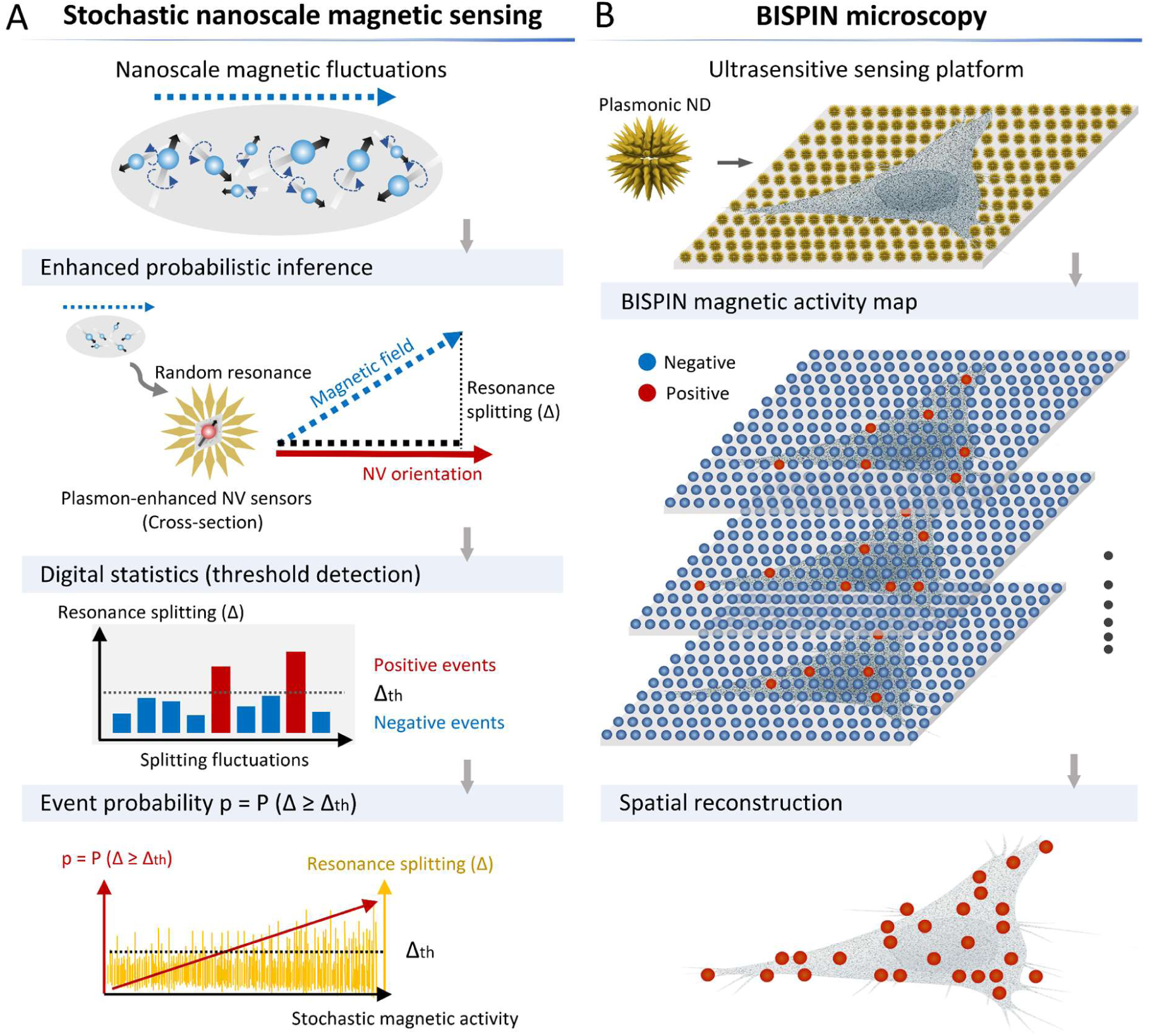
Conceptual framework of BISPIN microscopy. **(A)** Intrinsic cellular magnetic fluctuations originate from stochastic ionic transport and molecular currents. These magnetic perturbations are weak, randomly oriented, and temporally varying. NV magnetometry detects only the magnetic field projection along a fixed NV orientation, rendering analog estimation unstable under stochastic vector mixing. BISPIN introduces a predefined ODMR splitting threshold to classify magnetic perturbations into positive and negative digital events. Accumulated event statistics converge probabilistically to a descriptor of magnetic fluctuation strength within the measurement bandwidth. **(B)** BISPIN microscopy integrates plasmonic enhanced ND emitters into a high-density sensing substrate. Enhanced photon collection and ODMR contrast enable reliable digital event statistics. Spatial accumulation of positive events across the sensing plane yields stochastic reconstruction of intrinsic cellular magnetic fluctuation maps.

To address this limitation, we introduce BISPIN, a statistical framework that reformulates stochastic magnetometry in terms of probabilistic observables rather than deterministic magnetic values (Figure 1A). Instead of fitting instantaneous resonance shifts, BISPIN quantifies the probability that a magnetic perturbation produces a splitting excursion beyond a defined reference level. Reliable implementation of this probabilistic measurement requires high-fidelity detection of small resonance variations. We therefore employ plasmon-enhanced NV sensors that significantly increase photon collection efficiency and ODMR contrast (*22, 23*), thereby improving spin-readout fidelity in the weak-signal regime (Figure 1A, middle). The enhanced photon budget enables robust threshold classification of stochastic resonance fluctuations.

Operationally (Figure 1A, bottom), BISPIN defines a fixed ODMR splitting threshold (Δ_th_) determined from blank calibration measurements:

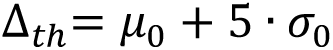

For each measurement, the observed splitting Δ is classified as a positive event if Δ≥ Δ_th_ and as a negative event otherwise. Repeated sampling over time yields an accumulated event probability (p):

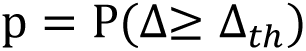

which serves as the primary digital observable.

Because stochastic magnetic perturbations broaden the distribution of Δ, the exceedance probability p increases monotonically with fluctuation strength within the experimental measurement bandwidth. For a broad class of unimodal fluctuation distributions, increasing variance at a fixed threshold leads to a higher exceedance probability, providing a statistical basis for using p as a descriptor of magnetic fluctuation strength. Since event classification depends only on whether a perturbation exceeds a fixed reference rather than on its precise instantaneous vector projection, BISPIN is intrinsically tolerant to random NV orientations, temporally stochastic magnetic dynamics, and biological heterogeneity. The probabilistic quantity p therefore provides a robust statistical descriptor of stochastic magnetic fluctuations without requiring reconstruction of a deterministic magnetic field.

Together, ultrasensitive NV nanosensors and probabilistic inference establish a digital magnetometry paradigm for stochastic nanoscale magnetic fluctuations. Rather than reconstructing deterministic magnetic fields, BISPIN extracts statistically reproducible information from stochastic magnetic disturbances and converts weak, fluctuating magnetic signals into a quantifiable probabilistic observable.

To extend this principle to imaging, BISPIN is implemented using a high-density array of plasmonic NDs (Figure 1B). Organized as a sensing array, these ultrabright NV emitters support wide-field acquisition of threshold-resolved magnetic events across the imaging plane. The resulting increase in photon statistics enables reliable event classification at each sensing site while maintaining nanoscale proximity to the sample, thereby allowing spatial mapping of stochastic magnetic activity.

Meanwhile, we apply this framework to controlled calibration measurements and subsequently to intrinsic magnetic activity at the cellular interface. In live cells, intrinsic cellular magnetic fluctuations arise from stochastic ionic transport and molecular current dynamics. These nanoscale magnetic perturbations are weak, rapidly varying, and randomly oriented within the cellular microenvironment. BISPIN microscopy is thereby increasing event fidelity while maintaining nanoscale proximity to living cells. High-density NV arrays form a sensing substrate that supports spatial accumulation of positive events, allowing stochastic reconstruction of cellular magnetic fluctuation maps. Accumulated positive-event distributions across the sensing plane yield a reconstructed representation of intrinsic cellular magnetic activity.

### 2.2 Photophysical properties of plasmon-enhanced NV sensors

To support reliable digital event statistics in BISPIN, we engineered an enclosed plasmonic ND architecture designed to enhance optical excitation and spin-dependent fluorescence readout of NV centers (Figure 2A). NV-hosting NDs (diameter ∼40 nm) were functionalized with single-stranded DNA through biotin-streptavidin linkages, while gold nanostars (Au NSs, diameter ∼90 nm) were modified with complementary DNA via Au-S bonds. DNA hybridization drove spontaneous self-assembly into a core-satellite configuration in which multiple Au NSs surround a central ND, forming an enclosed plasmonic nanocavity. Enhanced NV array substrates were then prepared by immobilizing enclosed plasmonic NDs on poly-L-lysine (PLL)-modified glass coverslips, followed by protective silica mineralization to stabilize the single NV layer while preserving its optical accessibility and sensing capability (Figure S1). Morphological characterizations via scanning electron microscopy (SEM) demonstrated that the average diameters of NDs, Au NSs, and enclosed plasmonic NDs were 43.7 nm, 96.2 nm, and 251.3 nm, respectively (Figure S2). Structural integrity of the core-satellite architecture was also confirmed experimentally by SEM (Figure 2A, insets i, ii, iii). Individual particles display complete enclosure of the central ND by Au NSs, and focused ion beam (FIB) cross-section imaging verified the formation of an enclosed nanocavity. Large-area SEM images demonstrated high assembly yield and structural reproducibility across arrays of particles.

**Figure 2.**
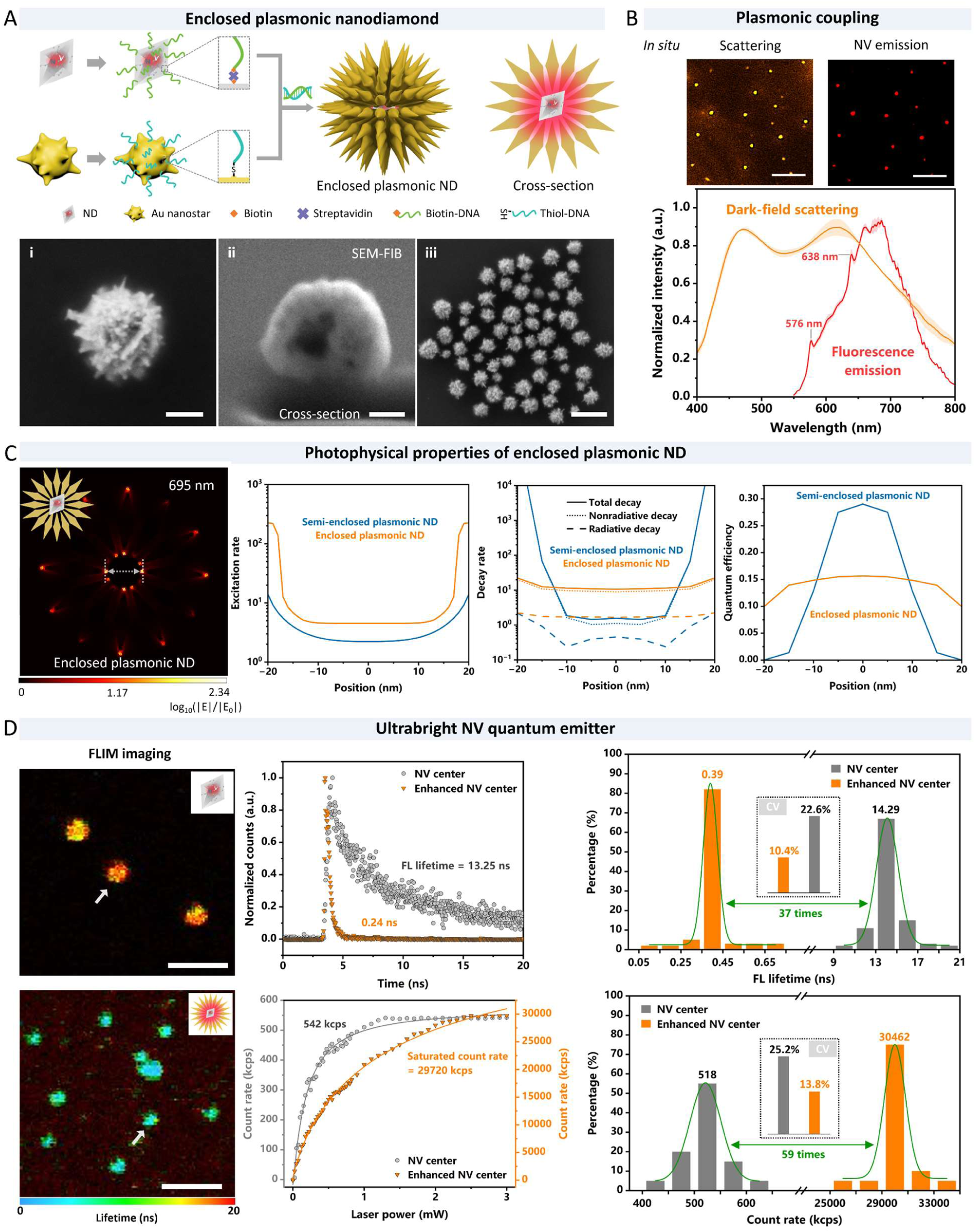
Photophysical characterization of enclosed plasmonic ND. **(A)** Schematic illustration of the DNA-guided self-assembly of enclosed plasmonic NDs. A 40 nm NV-hosting ND is surrounded by multiple 90 nm Au NSs through complementary DNA hybridization, forming an enclosed core-satellite nanocavity. Representative SEM image of a single enclosed plasmonic ND (inset i, scale bar: 100 nm), FIB cross-section image (inset ii, scale bar: 100 nm), and large-area SEM image of arrayed structures (inset iii, scale bar: 400 nm). **(B)** *In situ* optical verification of plasmon-NV coupling. Dark-field scattering image (left) and fluorescence emission image (right) of enclosed plasmonic NDs acquired from the same field of view (scale bar: 20 μm). The corresponding single-particle dark-field scattering spectrum and fluorescence emission spectrum are shown, revealing broadband plasmonic scattering (400-800 nm) overlapping with NV fluorescence (550-800 nm). Zero-phonon lines at 576 nm (NV^0^) and 638 nm (NV^-^) are preserved. Data are represented as mean ± s.d., where error bars represent the s.d. of n = 3 measurement replicates for each sample. **(C)** FDTD simulations of electromagnetic response in semi-enclosed (control) and enclosed plasmonic NDs. Left: electric field intensity enhancement at 695 nm under horizontal polarization. Middle: calculated excitation rate enhancement along the dashed line indicated in the field map. Right: calculated total (solid), radiative (dashed), and nonradiative (short-dashed) decay rate enhancements as a function of NV position. Position-dependent quantum efficiency derived from the radiative-to-total decay ratio is shown for both configurations. An intrinsic quantum efficiency of 0.7 was assumed for the dipole emitter. **(D)** Optical emission properties of NDs and enclosed plasmonic NDs. FLIM images (scale bar: 500 nm) and representative fluorescence decay curves show lifetime shortening in enclosed structures. Saturation curves of photon emission under increasing excitation power are shown with Logistic fitting. Statistical distributions of fluorescence lifetimes (n = 100 per group) and saturated count rates (n = 20 per group) are presented, with insets indicating CV and Gaussian fitting curves.

*In situ* optical characterization conducted on a custom-built optical system (set-up illustrated in Figure S3) further validated plasmon-emitter coupling (Figure 2B). Dark-field scattering imaging revealed the spatial distribution of Au NSs, while fluorescence imaging identified NV-hosting NDs within the same field of view. Strong spatial colocalization of scattering and fluorescence signals confirmed successful formation of the enclosed core-satellite nanostructure. Single-particle spectroscopy showed broadband plasmonic scattering extending from 400 to 800 nm, overlapping with the broad NV fluorescence emission spanning 550-800 nm. Distinct zero-phonon lines at 576 nm (NV^0^) and 638 nm (NV^-^) were clearly resolved (*24, 25*), indicating preservation of NV charge states within the plasmonic environment.

Finite-difference time-domain (FDTD) simulations were performed to evaluate the electromagnetic response of the enclosed nanocavity. The ND was modeled as a 40 nm sphere containing a single NV center represented as an electric dipole emitter with an intrinsic quantum efficiency of 0.7. For comparison, a semi-enclosed configuration consisting of a spherical ND surrounded by Au nanospheres was also simulated. At the principal NV emission wavelength (695 nm), the enclosed configuration exhibited substantially stronger localized electric field enhancement than the semi-enclosed control (Figure 2C, Figure S4). The most intense hotspots were concentrated at the Au NS-ND interfaces, reflecting enhanced near-field confinement within the nanocavity. Simulated excitation and decay-rate enhancements were evaluated as a function of emitter position inside the ND. In the enclosed geometry, both excitation and total decay rates remained significantly enhanced and comparatively insensitive to NV position, whereas the semi-enclosed configuration displayed pronounced spatial dependence with reduced enhancement toward the ND center. The enclosed nanocavity therefore mitigates sensitivity to the stochastic spatial distribution of NV centers within the ND. Calculated quantum efficiency remained stable across positions in the enclosed structure, indicating that radiative enhancement is maintained without excessive nonradiative loss.

Fluorescence lifetime imaging microscopy (FLIM) revealed pronounced lifetime shortening in enclosed plasmonic NDs compared to NV centers (Figure 2D). Representative decay curves showed a reduction from approximately 13 ns in NV centers to sub-nanosecond lifetimes in enhanced emitters. Statistical analysis of large-area FLIM images (Figure S5) across 100 particles yielded an average lifetime of 0.39 ns for enclosed structures, compared to 14.29 ns for NV centers. The reduced coefficient of variation (CV) observed for enhanced emitters indicates improved uniformity of optical response across particles. Photon emission saturation measurements further demonstrated substantial enhancement in emission rate. The average saturated count rate of enclosed structures exceeded that of NV centers by nearly two orders of magnitude, with reduced variability across particles. These results collectively indicate enhanced radiative decay and increased photon flux arising from the modified local density of optical states within the enclosed nanocavity.

The combined effects of increased excitation rate, accelerated radiative decay, and improved photon collection lead to significantly higher fluorescence brightness and improved ODMR contrast. For BISPIN, this enhancement directly improves digital threshold classification fidelity and accelerates statistical convergence of event probability measurements. The near position-independent enhancement within the enclosed nanocavity reduces variability associated with random NV location and orientation, thereby supporting robust stochastic magnetic sensing at the cellular interface.

### 2.3 Magnetic field measurement and resolution of plasmon-enhanced NV centers

To evaluate whether the enclosed plasmonic architecture improves magnetic-field measurement performance, optically detected magnetic resonance (ODMR) measurements were performed on NDs and enclosed plasmonic NDs under identical experimental conditions (Figure 3A). Samples were excited under wide-field total internal reflection fluorescence (TIRF) microscopy while a microwave antenna delivered frequency-swept microwave irradiation. The fabrication flowchart, structural parameter, and photograph of the omega-shaped microwave antenna are presented in Figures S6, S7, and S8, and a schematic of the microwave generating device is shown in Figure S9. A movable permanent magnet generated a tunable static magnetic field perpendicular to the substrate. ODMR spectra were acquired by scanning microwave frequency from 2820 to 2920 MHz with synchronized fluorescence readout at each frequency step.

**Figure 3.**
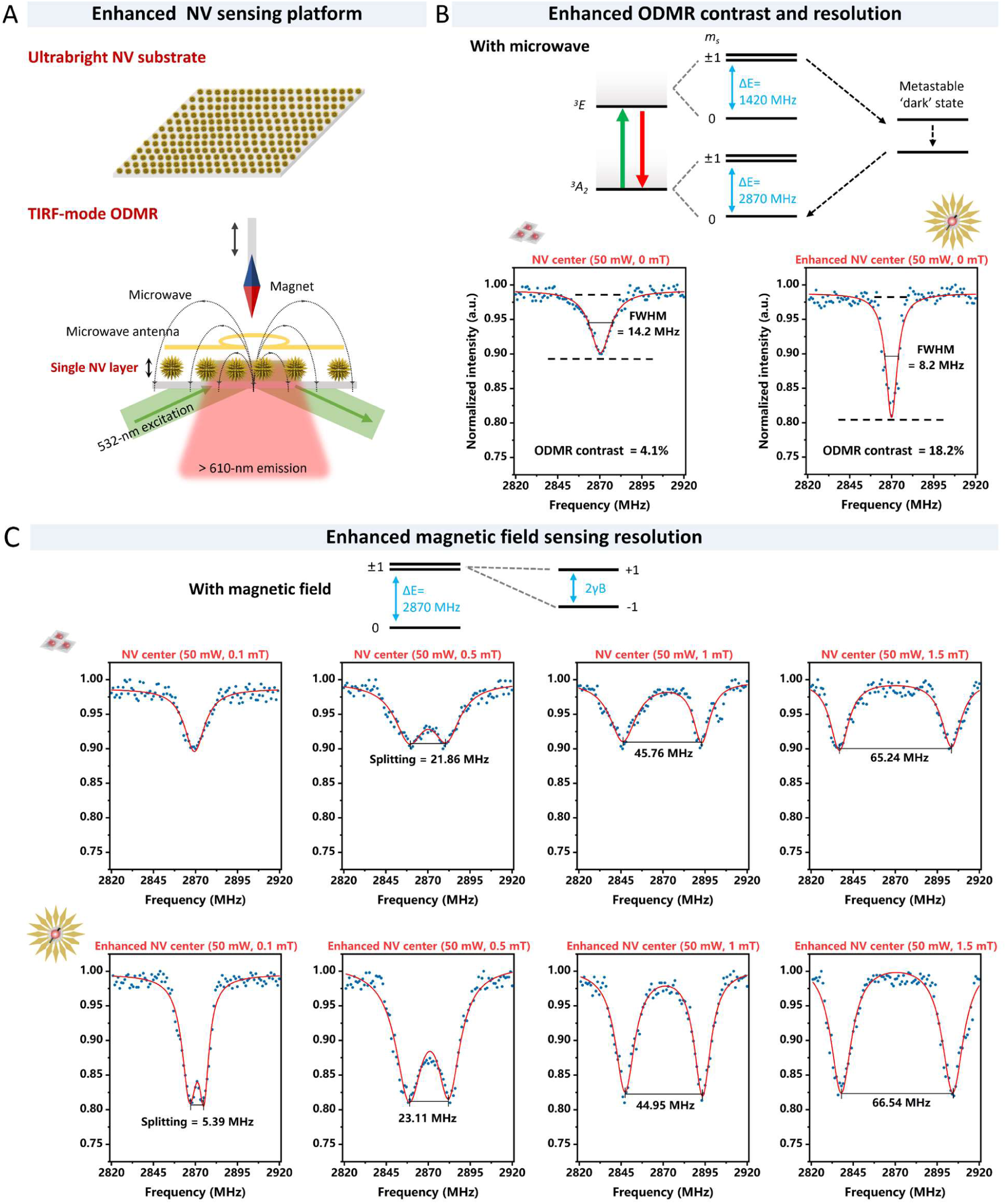
Magnetic field measurement and resolution of plasmon-enhanced NV centers. **(A)** Experimental configuration for wide-field ODMR measurements. An ultrabright plasmon-enhanced NV substrate is integrated with a microwave antenna for frequency-swept excitation, and a movable permanent magnet for applying a static magnetic field. Samples are excited at 532 nm under TIRF-mode illumination, and spin-dependent fluorescence emission (>610 nm) is collected for resonance analysis. **(B)** ODMR contrast and linewidth at zero magnetic field. Top: simplified NV ground-state energy level diagram illustrating microwave-driven transitions between *ms* = 0 and *ms* = ±1 sublevels and spin-dependent optical readout via intersystem crossing. Bottom: representative ODMR spectra of NDs and enclosed plasmonic NDs at 50 mW microwave power and 0 mT. Plasmon-enhanced structures exhibit increased contrast and reduced FWHM. Solid red curves represent Lorentzian fitting. **(C)** Magnetic-field-dependent ODMR measurements at a fixed microwave power of 50 mW. Resonance splitting induced by Zeeman interaction is shown for increasing static magnetic fields (0.1-1.5 mT) for NDs (top row) and enclosed plasmonic NDs (bottom row). Lorentzian fits are overlaid in red. Enhanced NDs retain resolvable resonance splitting at weak magnetic fields, where splitting in NDs becomes difficult to distinguish.

The underlying sensing principle follows the established NV center ground-state structure (*26*) (Figure 3B). In the absence of an external magnetic field, microwave excitation drives transitions between the *m_s_* = 0 and *m_s_* = ±1 spin sublevels near 2870 MHz. Under optical pumping, spin-dependent intersystem crossing to a metastable state produces reduced fluorescence for the *m_s_* = ±1 states, enabling optical readout of spin resonance. In the presence of a static magnetic field, Zeeman interaction lifts the degeneracy of the *m_s_* = ±1 states, resulting in two symmetrically split resonance frequencies. At zero magnetic field and fixed microwave power (50 mW), enclosed plasmonic NDs exhibit substantially increased ODMR contrast and reduced full width at half maximum (FWHM) compared with NDs (Figure 3B). Notably, across the full range of microwave powers tested (0-200 mW) under zero magnetic field, enclosed plasmonic NDs consistently display deeper resonance dips and narrower linewidths than NDs (Figure S10), indicating improved signal-to-noise ratio and spectral definition in spin-state readout. Quantitative analysis of spin-state readout performance—including ODMR contrast, FWHM, and derived magnetic sensitivity—is presented in Figure S11. Importantly, the resonance frequency remains centered near 2870 MHz, consistent with preservation of intrinsic NV spin properties.

Magnetic-field-dependent ODMR measurements further demonstrate improved spectral resolvability (Figure 3C). As the static magnetic field increases from 0.1 mT to 1.5 mT, Zeeman splitting of the *m_s_* = ±1 transitions increases linearly for both systems. However, at weak magnetic fields (e.g., 0.1 mT), splitting in NDs becomes difficult to resolve due to limited spectral contrast and broader linewidth, whereas enclosed plasmonic NDs retain clearly distinguishable dual resonance peaks. Lorentzian fitting confirms that resonance separation can be resolved at lower magnetic fields in the plasmon-enhanced structures. The improved magnetic-field resolution arises primarily from increased fluorescence brightness and enhanced ODMR contrast, which reduce uncertainty in determining resonance frequency minima. Narrower linewidths further improve the precision of frequency extraction. These combined effects enable more reliable detection of small resonance splittings under low magnetic fields.

For BISPIN, improved ODMR contrast and spectral resolution are critical because threshold classification relies on reproducible identification of resonance splittings relative to a predefined splitting reference. Enhanced photon flux and sharper spectral features increase the fidelity of digital event detection and reduce statistical ambiguity in determining whether a fluctuation exceeds the threshold. The enclosed plasmonic architecture therefore provides a stable magnetic-sensing platform that preserves intrinsic NV spin physics while improving magnetic-field resolvability within the biologically relevant low-field regime.

### 2.4 Ultrasensitive digital event statistics and validation of BISPIN quantification

To validate the quantitative behavior of BISPIN and benchmark the benefit of plasmon-enhanced NV emitters for digital event statistics, we first tested BISPIN under controlled static magnetic fields (Figure 4A). The detailed workflow of BISPIN, including thresholding and binary outcomes, is illustrated in Figure S12. ODMR-derived splitting metrics were converted into binary outcomes by applying a fixed threshold Δ_th_ defined from blank controls as Δ_th_= *μ*_0_ + 5 · *σ*_0_, where *μ*_0_ and *σ*_0_ denote the mean and standard deviation of the baseline splitting distribution. For each field strength, repeated measurements were acquired across multiple fields of view to accumulate event counts, and the positive-event fraction was computed as p = P(Δ≥ Δ_th_).

**Figure 4.**
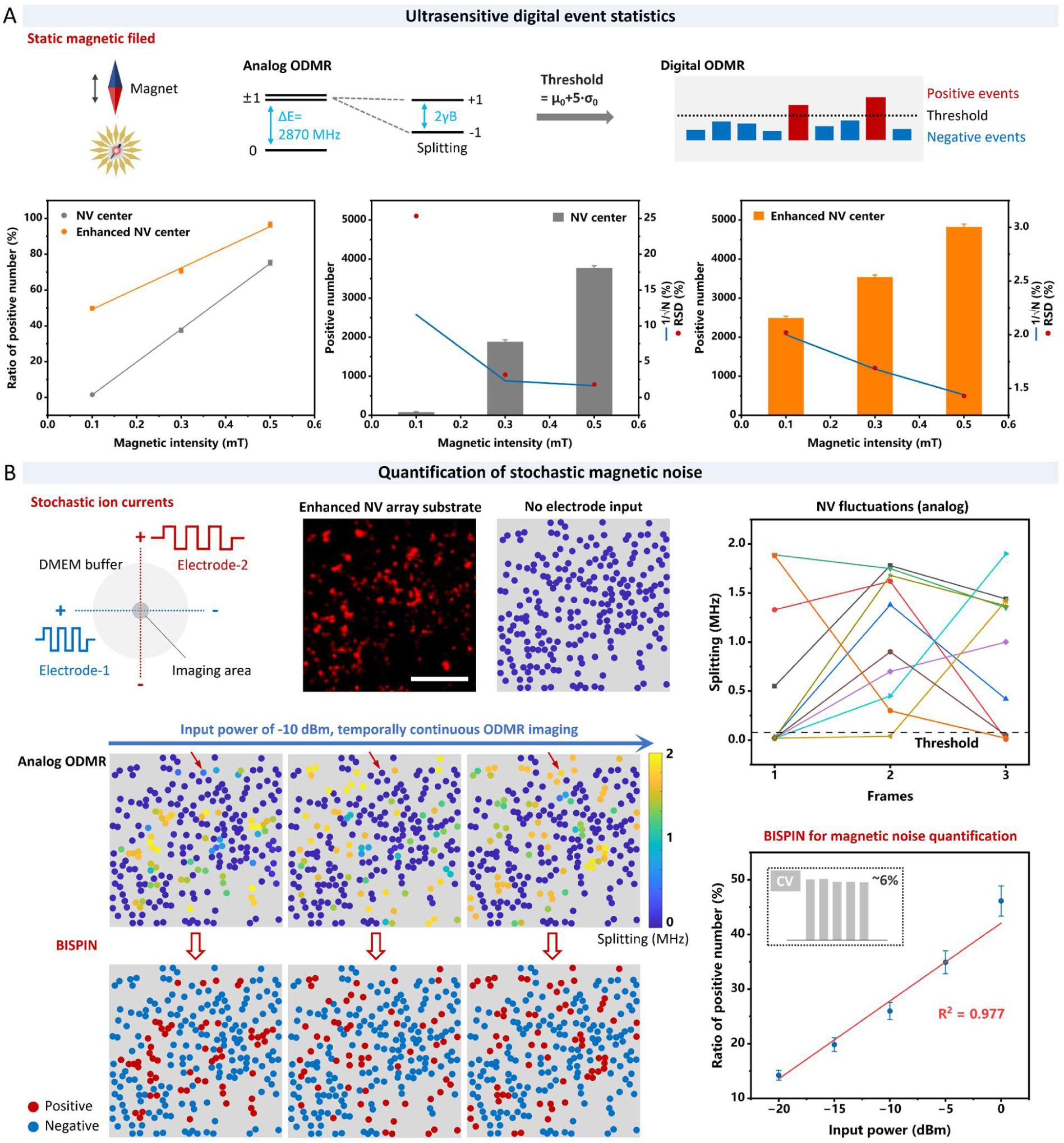
Ultrasensitive digital event statistics and stochastic magnetic-noise quantification using BISPIN. **(A)** Validation of digital threshold statistics under controlled static magnetic fields. Left: ratio of positive events as a function of applied magnetic field (0.1, 0.3, and 0.5 mT) for NDs and enclosed plasmonic NDs. Middle and right: total positive-event counts and corresponding RSD for NV substrates and enhanced NV substrates, respectively. ODMR-derived splitting metrics were converted into binary outcomes using a fixed threshold defined from baseline controls. Enhanced NV substrates yield higher positive-event fractions and reduced statistical variability, particularly in the weak-field regime. **(B)** Quantification of stochastic magnetic perturbations generated by ion-current excitation. Left (top): schematic of orthogonal electrode configuration in DMEM buffer for generating temporally varying ionic currents, and representative wide-field fluorescence image of enhanced NV array substrate (scale bar: 10 µm). Without electrode input, analog ODMR splitting maps establish the low-noise baseline for defining BISPIN threshold. Left (middle): analog ODMR splitting maps acquired under temporally continuous excitation (−10 dBm input power). Left (bottom): BISPIN converts these unstable analog readouts into binary outcomes at each pixel and time window, yielding binary event maps of positive and negative detections. Right (top): frame-to-frame analog splitting fluctuations illustrating stochastic variability relative to threshold. Right (bottom): BISPIN-based regional positive-event fractions as a function of electrode input power. The positive-event fraction exhibits a low CV (∼6%) and an approximately linear dependence on input power (R^2^ = 0.977), demonstrating a statistically convergent digital readout of stochastic magnetic disturbance strength. Data are represented as mean ± s.d., where error bars represent the s.d. of n = 3 measurement replicates for each sample.

At identical applied static fields (0.1, 0.3, and 0.5 mT), plasmon-enhanced NV substrates yielded substantially higher positive-event fractions than NV substrates (Figure 4A, bottom), indicating improved ability to resolve weak field-induced splitting excursions relative to the threshold. In parallel, the absolute number of positive events increased with magnetic field strength for both systems but was markedly larger for enhanced NV emitters (Figure 4A, bottom (left)), consistent with improved ODMR contrast and narrower linewidth (Figure 3) leading to more reliable classification of threshold exceedance. We next evaluated the statistical fidelity of the digital observable by analyzing the variability of positive-event counts. For a fixed acquisition size, the relative uncertainty in an event-counting process is expected to decrease with an increasing number of detected positive events. Consistent with this expectation, the enhanced NV platform exhibits reduced relative standard deviation (RSD) as magnetic field strength increases (Figure 4A, bottom (right)), approaching the behavior expected from counting statistics. In contrast, NV substrates show markedly elevated variability in the low-field regime, where positive events are sparse and analog ODMR fitting uncertainty more readily propagates into threshold classification (Figure 4A, bottom (middle)). These results indicate that plasmon-enhanced NV emitters improve not only the mean event response but also the stability and convergence of BISPIN event statistics in the weak-signal regime most relevant to cellular measurements.

Having established digital stability under static fields, we next validated BISPIN microscopy (set-up illustrated in Figure S13) for quantification of stochastic magnetic disturbances using a fully controlled extracellular calibration based on ion-current-driven magnetic noise (Figure 4B). Enhanced NV array substrates, which achieve ∼2 μm spatial resolution (Figure S14), were immersed in Dulbecco’s Modified Eagle Medium (DMEM) buffer and driven by orthogonal electrode pairs. Two alternating electrical signals with equal input power but incommensurate frequencies were applied to generate temporally varying ionic currents whose resultant magnetic perturbations fluctuate in magnitude and direction within the imaging region (set-up illustrated in Figure S15). Wide-field fluorescence imaging and ODMR measurements were conducted over a 30 μm^2^ area (Figure 4B, left (top)), approximating a single-cell-sized region. In the absence of electrode input, ODMR splitting maps remain near zero across the field of view, establishing a low-noise baseline from which the BISPIN threshold was defined. Under temporally continuous ODMR imaging at an input power of −10 dBm, conventional analog splitting maps exhibited strong frame-to-frame variability across pixels, reflecting the intrinsically stochastic and direction-mixed nature of the perturbations (Figure 4B, left (middle) and right (top)). BISPIN converts these unstable analog readouts into binary outcomes at each pixel and time window, yielding event maps of positive and negative detections (Figure 4B, left (bottom)).

Across the full range of input powers tested (−20, −15, −10, −5, and 0 dBm), conventional analog splitting maps consistently exhibited pronounced frame-to-frame variability across pixels, whereas BISPIN robustly yielded reliable positive- and negative-event maps (Figures S16-S20). Despite the random directionality and temporal variability of the ion-current-induced perturbations, the regional positive-event fraction increased reproducibly with increasing electrode input power (Figure 4B, right (bottom)). Across input powers, the BISPIN readout displayed low variability (CV = ∼6%) and an approximately linear dependence of positive-event fraction on input power (R^2^ = 0.977), demonstrating robust quantification of stochastic magnetic disturbance strength within a single-cell-sized field of view. Together, these benchmarks establish that BISPIN transforms unstable analog ODMR fluctuations into a statistically convergent digital observable, and that plasmon-enhanced NV emitters provide the event fidelity required for quantitative sensing in the weak and stochastic regime.

### 2.5 BISPIN microscopy for single-cell mapping of intrinsic stochastic magnetic activity

We next applied BISPIN microscopy to living cells cultured on the plasmon-enhanced NV sensing substrate to map intrinsic stochastic magnetic activity at the single-cell level (Figure 5A). Cells were placed in close proximity to a single NV sensing layer and imaged under TIRF-mode ODMR to maximize collection efficiency from near-interface NV emitters. For BISPIN analysis, ODMR-derived splitting metrics were converted into binary outcomes using a fixed threshold Δ_th_ defined from baseline controls, and positive-event fraction was used as the quantitative readout.

**Figure 5.**
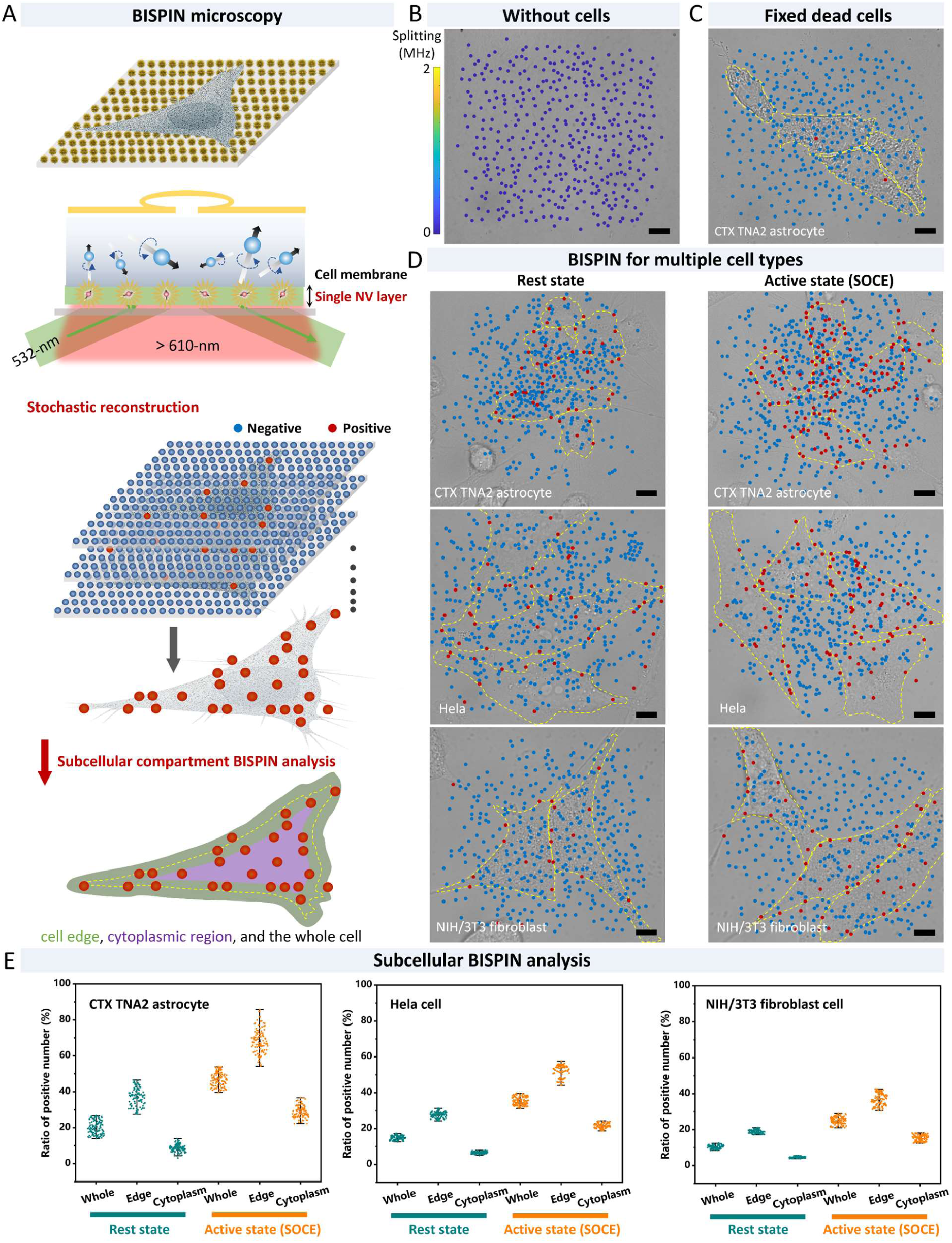
Single-cell BISPIN microscopy reveals state- and cell-type-dependent stochastic magnetic activity. **(A)** Schematic of BISPIN microscopy implemented on a plasmon-enhanced NV substrate for single-cell measurements. Cells are cultured directly above a single NV sensing layer and imaged under TIRF-mode ODMR. Analog splitting measurements are converted into binary classifications (positive/negative) using a fixed threshold, generating spatial event maps and compartment-resolved statistics. **(B)** Blank control measurement without cells. Wide-field analog splitting map remains near zero across the field of view (scale bar: 10 µm), establishing the low-noise baseline for defining BISPIN threshold. **(C)** Negative control with paraformaldehyde-fixed CTX TNA2 astrocytes. BISPIN event mapping reveals strongly reduced positive-event occurrence within the cell footprint (yellow dashed outline), indicating minimal contribution from static cellular components (scale bar: 10 µm). **(D)** BISPIN event maps of three cell types—CTX TNA2 astrocytes, Hela cells, and NIH/3T3 fibroblasts—under rest state and active state (SOCE). Positive (red) and negative (blue) detections are overlaid with cell boundaries (yellow dashed lines) (scale bars: 10 µm). SOCE stimulation increases the density of positive events in a cell-type-dependent manner. **(E)** Compartment-resolved BISPIN analysis of whole-cell, peripheral edge, and cytoplasmic regions for the three cell types (n = 100 per cell type) under rest state and active state (SOCE). Positive-event fractions (%) reveal reproducible differences between cell types and functional states.

We first established the baseline event distribution in the absence of cells (Figure 5B). Under identical imaging conditions, event maps showed predominantly negative classifications with minimal positive detections across the field of view, indicating low background exceedance under the defined Δ_th_. This blank control provides a reference level for thresholding and ensures that subsequent increases in positive-event fraction reflect cell-associated perturbations beyond instrumental background. To test whether passive cellular material could generate comparable exceedance events, we performed negative-control measurements using paraformaldehyde-fixed cells on the same substrate (Figure 5C). BISPIN analysis revealed strongly suppressed positive-event occurrence within the fixed-cell footprint relative to live-cell measurements, indicating that detectable events predominantly require active cellular processes rather than static cellular mass or surface attachment alone.

We then compared resting cells to a stimulated condition designed to upregulate transmembrane ionic transport. Store-operated calcium entry (SOCE) is a canonical pathway in which depletion of endoplasmic reticulum Ca^2+^ stores triggers extracellular Ca^2+^ influx through plasma-membrane channels (*27, 28*), increasing membrane current activity and associated electrodynamic fluctuations. Fluctuating transmembrane currents generate magnetic fields proportional to instantaneous current amplitude according to the Biot-Savart relation. In our experiments, SOCE stimulation—achieved via thapsigargin-induced store depletion followed by elevated extracellular Ca^2+^ (validated by Fluo-4 AM; Figure S21)—produced a clear increase in BISPIN positive-event density across cell types (Figure 5D). The enhancement was most prominent near the cell periphery, consistent with SOCE being mediated by membrane-localized ion channels and thus expected to increase near-membrane electrodynamic activity sensed by the shallow NV layer.

We performed BISPIN microscopy across three cell types—CTX TNA2 astrocytes, Hela cells, and NIH/3T3 fibroblasts—under rest state and active state (SOCE) (Figure 5D). Schematic of BISPIN thresholding for quantifying cellular stochastic magnetism is illustrated in Figure S22. In each case, cell outlines were defined from optical images and overlaid with BISPIN event maps, in which positive detections (red) and negative detections (blue) report the spatial distribution of threshold exceedances accumulated over the acquisition window. Compared with rest state, active state (SOCE) increased the positive-event fraction in all three cell types, yielding reproducible cell-state differences. Importantly, in cases where cellular magnetic fluctuations are weak and yield excessively low positive-event fractions, statistical significance can be improved by temporally accumulating binary-event frames, increasing the effective sampling of stochastic events (Figure S23). This frame-accumulation strategy enhances the reliability of the calculated positive-event fractions in reflecting underlying cellular magnetic activity, enabling meaningful BISPIN analysis even under low-signal conditions.

To quantify spatial heterogeneity, we performed subcellular BISPIN analysis by segmenting each cell into the whole-cell region, a peripheral edge compartment, and an interior cytoplasmic compartment (Figures 5A and E). Schematic of subcellular compartment segmentation is illustrated in Figure S24. Cell boundaries are defined from bright-field images, with a 6 μm-wide peripheral edge (±3 μm from the boundary) spanning extracellular and intracellular sides, while regions located more than 3 μm inside the boundary were defined as interior cytoplasmic regions. Across cell types, astrocytes displayed the highest positive-event fractions and the strongest edge-enriched response under stimulation, Hela cells showed intermediate responses with clear state dependence, and NIH/3T3 fibroblasts exhibited comparatively lower and more uniform event fractions. These subcellular-level profiles constitute reproducible single-cell fingerprints of stochastic magnetic activity.

The BISPIN readout reports stochastic magnetic disturbances within the experimental bandwidth and within the sensing volume set by NV depth and optical collection geometry. A plausible dominant contributor in living cells is ionic current activity associated with membrane transport processes, because fluctuating charge motion generates magnetic fields and SOCE stimulation selectively increases channel-mediated ionic flux. Additional contributors may include stochastic intracellular currents and charge redistribution driven by organelle activity and metabolic processes. While BISPIN does not assign a unique microscopic source at each pixel, the live-versus-fixed contrast and SOCE-dependent modulation support an interpretation that the measured signal primarily reflects activity-dependent electrodynamic fluctuations rather than static magnetic impurities or passive cellular material.

Together, these results demonstrate that BISPIN microscopy enables single-cell mapping of intrinsic stochastic magnetic activity using a fixed digital threshold and event statistics. By converting temporally variable ODMR signals into spatially resolved, statistically convergent exceedance maps, BISPIN provides a robust label-free and contact-free approach for comparing cell types, functional states, and subcellular compartments in the weak and stochastic regime.

### 2.6 Machine-learning-assisted decoding of single-cell bio-spin signatures

To evaluate whether BISPIN-derived digital event statistics encode sufficient information to distinguish cellular identity and functional state, we applied supervised machine learning to single-cell bio-spin signatures extracted from whole-cell measurements (Figure 6A). For each cell, the compartment-resolved positive-event fractions obtained from BISPIN analysis were used as the primary quantitative descriptor of stochastic magnetic activity. These values served as input features for classification across six experimental groups: CTX TNA2 astrocytes, Hela cells, and NIH/3T3 fibroblasts under rest state and active state (SOCE).

**Figure 6.**
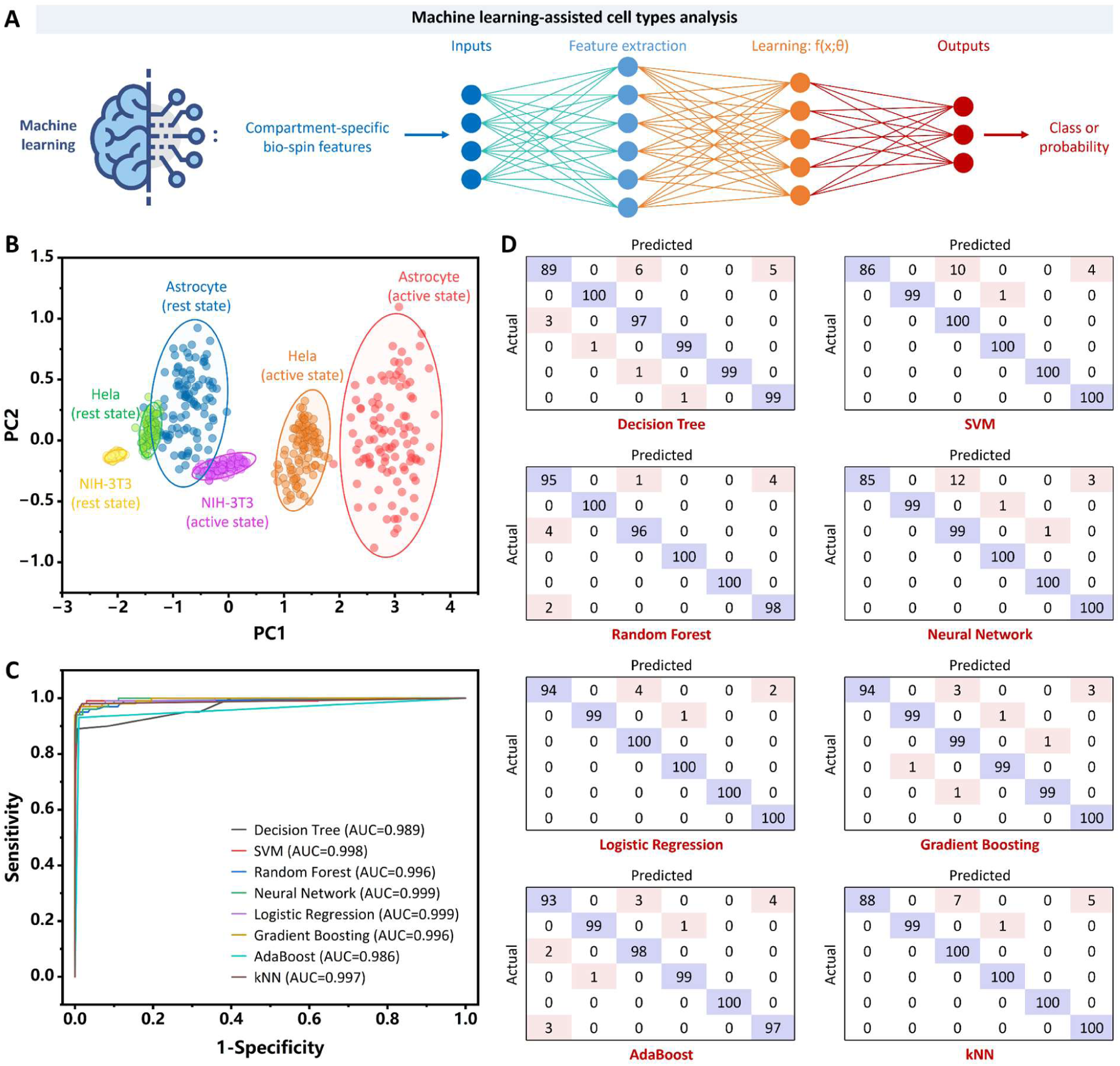
Machine-learning-assisted classification of single-cell bio-spin signatures. **(A)** Workflow integrating BISPIN-derived compartment-resolved positive-event fractions with supervised machine learning. Single-cell bio-spin signatures are used as input features to train classification models that assign samples to cell-type and functional-state classes. **(B)** PCA of six experimental groups—CTX TNA2 astrocytes, Hela cells, and NIH/3T3 fibroblasts under rest state and active state (SOCE). Distinct clustering indicates structured separation in feature space. **(C)** ROC curves for eight representative classification algorithms. AUC values range from 0.986 to 0.999, demonstrating robust class separability. **(D)** Confusion matrices showing classification performance across the six classes. Low misclassification rates indicate accurate decoding of cell identity and functional state from BISPIN-derived bio-spin statistics.

We first visualized the intrinsic structure of the dataset using principal component analysis (PCA) (Figure 6B). Projection onto the first two principal components revealed clear clustering by both cell type and functional state. Even without nonlinear transformation, the six groups occupied distinct regions in feature space, indicating that compartment-resolved bio-spin signatures exhibit structured variability rather than random dispersion.

To quantify separability, we implemented eight representative machine learning models spanning linear (Logistic Regression), kernel-based (SVM), tree-based (Decision Tree), ensemble (Random Forest, Gradient Boosting, AdaBoost), Neural Network, and instance-based (kNN) approaches. Receiver operating characteristic (ROC) analysis demonstrated consistently high discriminative performance across models (Figure 6C), with area under the curve (AUC) values ranging from 0.986 to 0.999. Importantly, linear and nonlinear models achieved comparable accuracy, indicating that class separation does not rely on complex nonlinear decision boundaries but instead reflects structured differences in bio-spin statistics. Confusion matrix analysis (Figure 6D) confirmed low misclassification rates across all six classes, supporting the reproducibility of the measured signatures. Misclassifications were sparse and distributed rather than concentrated in specific group pairs, suggesting that classification performance is not driven by trivial pairwise separability.

Together, these results demonstrate that stochastic magnetic activity quantified by BISPIN forms reproducible single-cell signatures capable of distinguishing both cellular identity and functional state. Without relying on exogenous labels or predefined molecular markers, BISPIN-derived bio-spin signatures provide an orthogonal phenotypic dimension reflecting intrinsic electrodynamic activity in living cells.

## 3. Conclusions

In this work, we established a digital statistical framework—BISPIN—for quantitative access to intrinsic stochastic magnetic activity in living cells. Rather than attempting to reconstruct unstable instantaneous magnetic amplitudes, BISPIN reformulates nanoscale magnetometry as an event-based inference problem. By introducing a fixed threshold in spin-resolved optical readout and converting analog fluctuations into statistically convergent digital observables, BISPIN enables robust quantification of weak, randomly oriented, and temporally varying magnetic disturbances. Coupled with plasmon-enhanced NV sensing substrates, this framework provides improved spin-readout fidelity, enhanced spectral resolution, and stable event statistics in the low-signal regime relevant to biological systems. Using this approach, we demonstrated reproducible mapping of stochastic magnetic activity at the single-cell level, discrimination of cell types and functional states, and data-driven classification of bio-spin signatures without electrical contact, molecular labeling, or exogenous reporters.

Conceptually, BISPIN shifts the paradigm of quantum biosensing from deterministic field reconstruction toward statistical characterization of stochastic electrodynamic fluctuations. This shift parallels the transition from deterministic trajectory tracking to probabilistic inference in modern statistical physics and information theory. In living systems, magnetic signals are inherently random, heterogeneous, and bandwidth-limited. By embracing stochasticity as an informative signal dimension rather than treating it as measurement noise, BISPIN establishes a measurement principle that is inherently aligned with the physics of biological dynamics. Importantly, the digital inference strategy introduced here is not restricted to ODMR-based splitting measurements. NV centers provide multiple complementary observables—including longitudinal relaxation time (T_1_) (*29, 30*), transverse coherence time (T_2_) (*31*), inhomogeneous dephasing time (T_2_*) (*32*), and other spin-dependent contrasts—each sensitive to distinct spectral components of magnetic noise. Extending digital threshold-based probabilistic inference to these modalities could enable bandwidth-resolved and mechanism-sensitive bio-spin microscopy, in which different spin parameters report on different classes of stochastic electrodynamic processes. Such an approach would expand BISPIN from a single-contrast method into a multidimensional quantum sensing framework.

More broadly, this work lays the foundation for a quantitative electrodynamic dimension of cellular phenotyping. Just as fluorescence microscopy enabled molecular mapping and single-molecule localization transformed spatial resolution, probabilistic spin-based inference opens a route to statistically resolvable magnetic signatures in living matter. Future developments integrating advanced NV sensing protocols, engineered spin coherence control, and scalable digital inference may enable comprehensive characterization of stochastic bio-spin activity across spatial scales, cell types, and physiological conditions. By transforming intrinsically fluctuating magnetic activity into a measurable and classifiable physical observable, BISPIN provides a general framework for probing electrodynamic processes in living systems and establishes a platform upon which future quantum biosensing methodologies can be built.

## Associated content

Supporting information: Sections 1-10; Figures S1-S24.

## Author Contributions

Le Liang conceived and mentored this research. Weiming Lin and Tao Ding contributed equally to this work. All authors edited the manuscript and approved the final version.

## Declaration of competing interest

The authors have no known competing financial interests or personal relationships that could have appeared to influence the work reported in this paper.

## Acknowledgements

We would like to acknowledge the support from the National Natural Science Foundation of China (Grant No. 22374112), the Fundamental Research Funds for the Central Universities, and the Start-Up Fund of Wuhan University. We also thank the Core Facility of Wuhan University for providing access to SEM and FLIM instruments.

## Supporting information

### Section 1: Chemicals and materials

Hydrogen tetrachloroaurate (HAuCl_4_·3H_2_O, 99%) and Ascorbic acid (AA, 99%) were purchased from Meryer Biochemical Technology (China). Silver nitrate (AgNO_3_) was purchased from Chron Chemicals (China). Poly dimethyl diallyl ammonium chloride (PDDA) was purchased from Heowns Biochem LLC (China). Sodium borohydride (NaBH_4_, 98%), sodium citrate (Na_3_C_6_H_5_O_7_·2H_2_O, 98%), hydrogen peroxide (H_2_O_2_, 30%), tris(2-carboxyethyl)-phosphine (TCEP), 3-(trimethoxysilyl)propyltrimethylammonium chloride (TMAPS), and tetraethyl orthosilicate (TEOS) were obtained from Aladdin Biochemical Technology (China). Sulfuric acid (H_2_SO_4_) was purchased from Sinopharm Chemical Reagent (China). DNA strands were synthesized by Sangon Biotech (China) and stored at −20 °C. Amicon Ultra-0.5 mL centrifugal filters and 0.22 μm membrane filters were purchased from Merck Millipore (USA). Biotinylated fluorescent nanodiamonds (NDs) (15 nM, 40 nm in diameter) were purchased from Adámas Nanotechnologies (USA). Sodium chloride (NaCl) and calcium chloride (CaCl_2_) were purchased from Energy Chemical (China). Tris-acetate-EDTA (TAE) buffer and Tris-borate-EDTA (TBE) buffer were purchased from Macklin Biochemical Technology (China). Cells (CTX TNA2 astrocytes, Hela, and NIH/3T3 fibroblasts) were obtained from the American Type Culture Collection. Dulbecco’s Modified Eagle Medium (DMEM), fetal bovine serum (FBS), and penicillin-streptomycin solution were purchased from Gibco (USA). Dimethyl sulfoxide (DMSO) and poly-L-lysine (PLL, molecular weight 150000-300000) were purchased from Sigma-Aldrich (USA). Ca^2+^-free Tyrode buffer was purchased from Yuanye Biotechnology (China). Thapsigargin in DMSO solution were purchased from Abcam (UK). Fluo-4 acetoxymethyl ester (Fluo-4 AM) was purchased from Thermo Fisher Scientific (USA). Ultrapure water (18.2 MΩ·cm at 25 °C) produced by a Millipore purification system (Milli-Q IQ Element, Merck, USA) was used throughout our experiments.

### Section 2: Preparation of gold nanostar (Au NS)-DNA conjugates

#### Supplementary Note S1: Synthesis of 13 nm gold nanoparticle seeds

To prepare the seed solution, an aqueous HAuCl_4_·3H_2_O solution was initially diluted with ultrapure water under continuous magnetic stirring, which was maintained throughout the entire process. Subsequently, sodium citrate was injected into the mixture. A freshly prepared NaBH_4_ solution was then introduced at an extremely slow rate (one drop every 5 s). Finally, the reaction mixture was left to stir overnight at room temperature.

**Supplementary Table S1:** Recipe for synthesis of 13 nm gold nanoparticle seeds.

|  | Final desired concentration | Stock concentration | To add |
| --- | --- | --- | --- |
| $\text{H}_2\text{O}$ | - | - | 90 mL |
| $\text{HAuCl}_4 \cdot 3\text{H}_2\text{O}$ | 0.299 mM | 28.4 mM | 1 mL |
| $\text{Na}_3\text{C}_6\text{H}_5\text{O}_7 \cdot 2\text{H}_2\text{O}$ | 0.817 mM | 38.8 mM | 2 mL |
| $\text{NaBH}_4$ | 0.000789 wt % | 0.075% wt % in 38.8 mM<br>$\text{Na}_3\text{C}_6\text{H}_5\text{O}_7 \cdot 2\text{H}_2\text{O}$ solution | 1 mL |
| Total | - | - | 94 mL |

#### Supplementary Note S2: Synthesis of Au NSs

Au NSs were synthesized via a seed-mediated growth approach. Initially, the pre-synthesized 13 nm gold nanoparticle seeds were dispersed in a 10-fold diluted PDDA solution. Separately, the growth solution was prepared by sequentially introducing HAuCl_4_, AgNO_3_, and AA into a PDDA solution under continuous and vigorous stirring. To trigger the Au NS synthesis, the diluted seed solution was added to the growth mixture. The reaction was then allowed to proceed under vigorous stirring overnight at room temperature. Finally, the obtained Au NS dispersion was centrifuged to achieve a 10-fold concentration.

**Supplementary Table S2:** Recipe for diluting 13 nm gold nanoparticle seeds.

|  | Final desired concentration | Stock concentration | To add |
| --- | --- | --- | --- |
| PDDA solution | 100 mg/mL | 111 mg/mL | 9 mL |
| 13 nm gold nanoparticle seeds | 0.452 nM | 4.52 nM | 1 mL |
| Total | - | - | 10 mL |

**Supplementary Table S3:** Recipe for synthesis of Au NSs.

|  | Final desired concentration | Stock concentration | To add |
| --- | --- | --- | --- |
| PDDA solution | 89.3 mg/mL | 100 mg/mL | 10 mL |
| HAuCl<sub>4</sub>·3H<sub>2</sub>O | 0.223 mM | 5 mM | 500 µL |
| AgNO<sub>3</sub> | 0.357 mM | 8 mM | 500 µL |
| AA | 0.893 mM | 100 mM | 100 µL |
| Diluted 13 nm gold nanoparticle seeds solution | 0.00404 nM | 0.452 nM | 100 µL |
| Total | - | - | 11.2 mL |

#### Supplementary Note S3: Synthesis of Au NS-DNA conjugates

Thiolated DNA strands were initially reduced using TCEP in an aqueous solution. Subsequently, the reduced DNA strands were incubated with Au NSs at a 50000:1 molar ratio in a NaCl-containing TBE buffer. This mixture was maintained at room temperature for 10 h under constant agitation at 400 rpm. Finally, to eliminate unbound DNA, the conjugate solution underwent a minimum of three washing cycles with 1×TAE buffer via centrifugation (3500 rpm, 3 min per cycle) using Amicon Ultra-0.5 mL filters (100 kDa MWCO).

**Supplementary Table S4:** Recipe for synthesis of Au NS-DNA conjugates.

|  | Final desired concentration | Stock concentration | To add |
| --- | --- | --- | --- |
| Thiolated DNA | 0.75 $\mu\text{M}$ | 250 $\mu\text{M}$ | 1.5 $\mu\text{L}$ |
| TCEP | 0.15 mM | 10 mM | 7.5 $\mu\text{L}$ |
| Au NSs solution | 15.0 pM | 97.6 pM | 76.8 $\mu\text{L}$ |
| TBE | 0.5× | 5× | 50 $\mu\text{L}$ |
| NaCl | 50 mM | 5000 mM | 5 $\mu\text{L}$ |
| Total | - | - | 500 $\mu\text{L}$ |

### Section 3: Preparation of enhanced NV array substrate

#### Supplementary Note S4: Preparation of ND-DNA conjugates

Prior to functionalization, a 1 mg/mL colloidal dispersion of 40 nm NDs was mixed with freshly dissolved streptavidin in an aqueous solution. This mixture was incubated at room temperature for 2 h under constant agitation at 400 rpm. To eliminate unbound streptavidin, the resulting ND-avidin conjugates were purified via three cycles of centrifugation at 5000 rpm for 10 min. Subsequently, the purified conjugates were incubated with biotinylated DNA at room temperature for an additional 2 h under constant agitation at 400 rpm. The final ND-DNA conjugates were separated from excess biotin-DNA through an identical three-step centrifugation process (5000 rpm, 10 min).

**Supplementary Table S5:** Recipe for synthesis of ND-DNA conjugates.

|  | Final desired concentration | Stock concentration | To add |
| --- | --- | --- | --- |
| Biotin DNA modified ND | 0.02 mg/mL | 1 mg/mL | 2 μL |
| Thiol DNA modified Au NS | 0.025 mg/mL | 1 mg/mL | 2.5 μL |
| Biotin-DNA | 0.0045 μM | 100 μM | 4.5 μL |
| Final | - | - | 100 μL |

#### Supplementary Note S5: Synthesis of enclosed plasmonic NDs

To fabricate enclosed plasmonic NDs, DNA-functionalized NDs were mixed with DNA-modified Au NSs at a predetermined ratio in TAE buffer. This mixture was incubated for 1 h to facilitate DNA hybridization. The resulting enclosed plasmonic NDs were then purified via centrifugation at 1700 rpm for 3 min. DNA sequences used in this work were listed below (5’ to 3’).

Au NS attachment: ATGATATAGACGTTGTGGCAAAAAAAA-thiol

ND attachment: GCCACAACGTCTATATCATAAAAAAAA-biotin

**Supplementary Table S6:** Recipe for synthesis of enclosed plasmonic NDs.

|  | Final desired concentration | Stock concentration | To add |
| --- | --- | --- | --- |
| Biotin DNA modified ND | 0.0008 mg/mL | 0.02 mg/mL | 20 μL |
| Thiol DNA modified Au NS | 15 pM | 75 pM | 100 μL |
| TAE buffer | 1× | 1.32× | 380 μL |
| Total | - | - | 500 μL |

#### Supplementary Note S6: Preparation of enhanced NV array substrate

A Glass coverslip was treated by oxygen plasma (180 W, 45 s) to activate the surface. A 100 μL aliquot of PLL (molecular weight 150000-300000) was added to the coverslip and incubated for 10 min to form a positively charged adhesion layer. The solution was then removed, and the surface was allowed to air-dry naturally. Next, 30 μL of the enclosed plasmonic ND stock suspension (6 pM) was deposited onto the PLL-treated surface and incubated for 30 min to allow particle adsorption. The coverslip was subsequently rinsed with ultrapure water to remove unbound particles and air-dried naturally.

To form a protective mineralized layer on the substrate, a silica mineralization precursor solution was first prepared. Specifically, 400 μL of TMAPS was slowly added to 20 mL of 1×TAE buffer in 20 μL increments with 5 s intervals between additions under continuous stirring. The mixture was stirred for 20 min. Subsequently, 400 μL of TEOS was added in the same incremental manner (20 μL per addition with 5 s intervals), followed by an additional 20 min of stirring to obtain the mineralization precursor solution. A 1 mL aliquot of the precursor solution was then added to the enclosed plasmonic ND-coated coverslip, and the substrate was incubated under static conditions for 48 h to allow formation of the mineralized layer. After incubation, the mineralization solution was removed, and the substrate was sequentially rinsed with 1 mL of anhydrous ethanol and ultrapure water while gently shaking the coverslip to remove residual reagents. The liquid was then removed and the substrate was air-dried, yielding the final enhanced NV array substrate.

**Figure S1.**
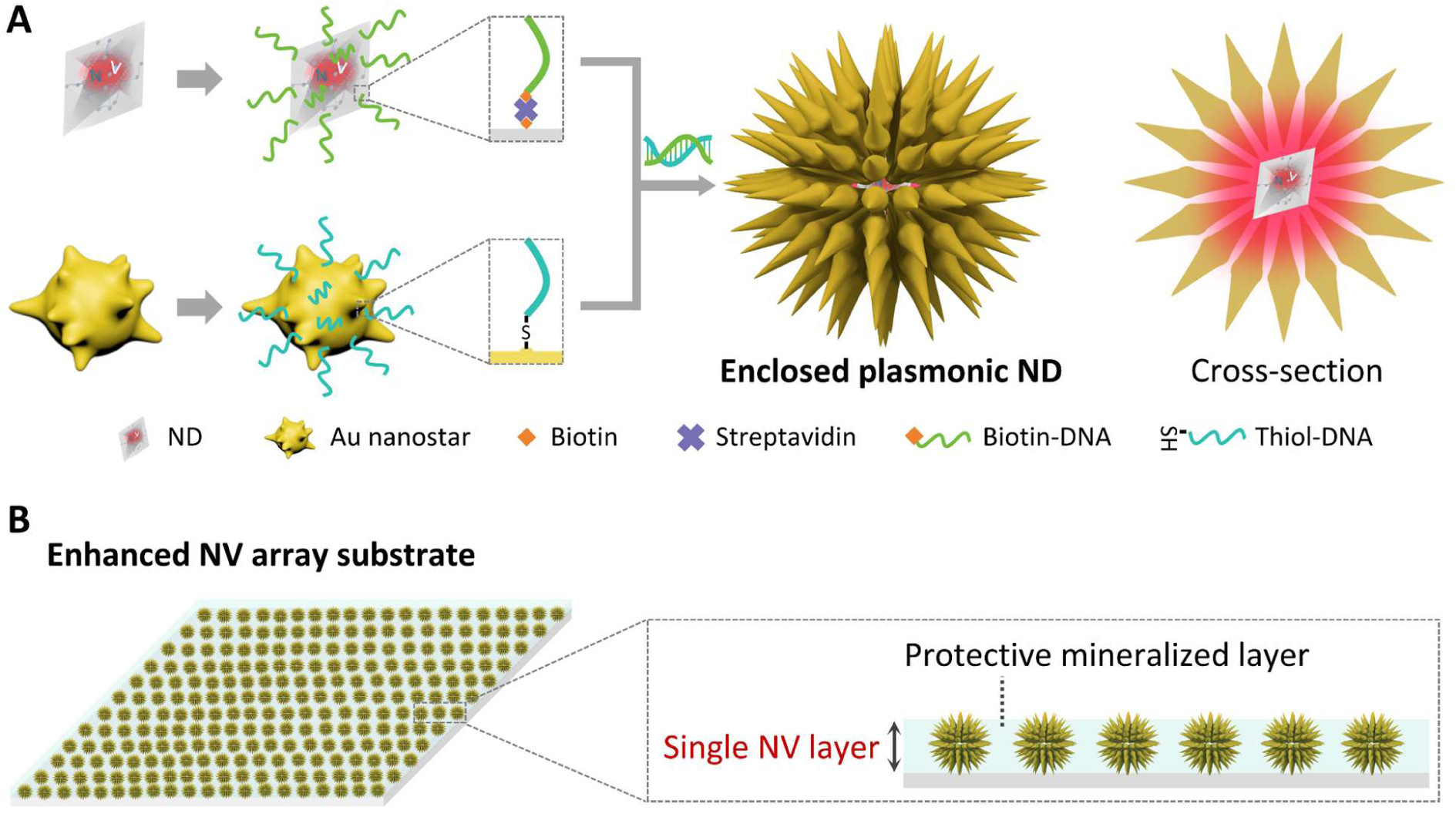
Preparation of enhanced NV array substrate. **(A)** Schematic illustration of the DNA-guided self-assembly of enclosed plasmonic NDs. A 40 nm NV-hosting ND is surrounded by multiple 90 nm Au NSs through complementary DNA hybridization, forming an enclosed core-satellite nanocavity. **(B)** A PLL-modified glass coverslip was used to immobilize enclosed plasmonic NDs, forming an enhanced NV array substrate. The substrate was subsequently subjected to silica mineralization via TMAPS-TEOS precursors, generating a thin protective layer that stabilizes the single NV layer while preserving its optical accessibility and sensing capability.

### Section 4: Morphological characterization

#### Supplementary Note S7: Scanning electron microscopy (SEM) imaging

Prior to sample deposition, silicon wafers were cleaned by immersion in a piranha solution for 20 min, followed by ultrasonication in ultrapure water for 20 min, and subsequently dried under a stream of compressed nitrogen. A 10 μL droplet of the sample was then cast onto the cleaned wafer and allowed to adsorb for 10 min. After rinsing the surface with ultrapure water, the substrate was dried at 25 °C. Morphological characterization was performed using a Field Emission SEM (Zeiss GeminiSEM 500, Germany) operating at an acceleration voltage of 5 kV.

#### Supplementary Note S8: Focused ion beam (FIB)-SEM imaging

To investigate the internal morphology of the capsids, the samples were analyzed using a FIB-SEM system. Prior to milling, the specimens were sputter-coated with a thin gold layer to enhance electrical conductivity and mitigate charging effects during imaging. A Ga ion beam was then employed to mill the sample and expose its cross-section. Finally, the internal structures were imaged utilizing a FIB SEM (TESCAN SOLARIS, Czech Republic) operating at an accelerating voltage of 3 keV.

#### Supplementary Note S9: Particle size analysis based on SEM imaging

The particle size distributions of samples were quantified from SEM images. For each sample type, SEM micrographs were acquired under identical imaging conditions to ensure comparability. Individual particle diameters were measured using the ROI Manager tool in Fiji (v2.9.0 or later). In each group, 100 particles were randomly selected and their diameters were recorded. The collected data were subsequently used to generate particle size histograms, from which the mean diameter and overall distribution profiles were extracted. All image analyses were performed in a blinded manner to minimize selection bias.

**Figure S2.**
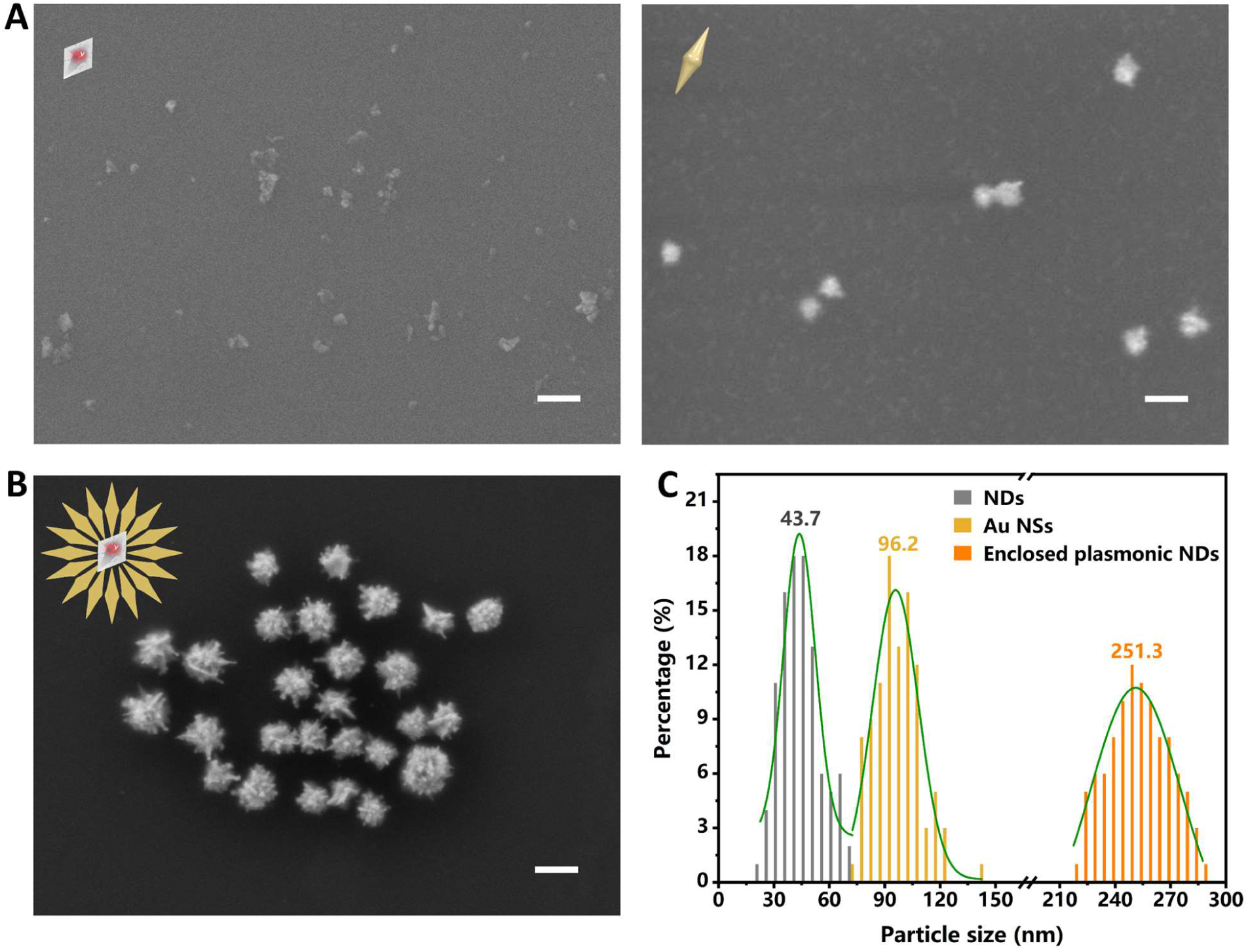
Morphological characterization of NDs, Au NSs, and enclosed plasmonic NDs. Representative large-area SEM images (scale bar: 200 nm) of **(A)** NDs, Au NSs, and **(B)** enclosed plasmonic NDs. **(C)** Statistical distributions of particle size (n = 100 per group) were obtained from large-area SEM images. Curves represent Gaussian fitting. The average diameters of NDs, Au NSs, and enclosed plasmonic NDs were 43.7 nm, 96.2 nm, and 251.3 nm, respectively.

### Section 5: Photophysical properties characterization

#### Supplementary Note S10: *In situ* dark-field and fluorescence imaging with single-particle spectroscopy

*In situ* optical measurements were performed at room temperature under ambient conditions using a custom-built wide-field imaging and single particle spectroscopy system.

Dark-field scattering was characterized on an inverted microscope (Nikon, Japan) equipped with a dark-field condenser (NA=1.0) and a 40× air objective (NA=0.6). The sample was illuminated by a halogen lamp (white light), and the scattered light was captured by a true-color camera for dark-field imaging.

Fluorescence imaging was performed using a 532 nm laser (OBIS Laser Box, Coherent Inc., USA) filtered through a 532±2 nm band-pass filter and reflected by a 532 nm long-pass dichroic mirror. The beam was focused onto the sample through the same 40× objective (NA=0.6). Emitted photoluminescence (PL) was collected by the objective, separated from the excitation laser by the dichroic mirror, passed through a 610 nm long-pass filter to suppress residual scattering, and recorded by a back-illuminated sCMOS camera (KURO 1200B, Princeton Instruments, USA).

For single-particle spectroscopy, the PL from a manually defined region of interest (ROI) under the same excitation configuration was further filtered by a 537 nm long-pass filter and directed to a grating spectrometer (SpectraPro HRS-500, Princeton Instruments, USA) equipped with a 600 g/mm grating and a back-illuminated sCMOS camera (KURO 1200B, Princeton Instruments, USA) for scattering and fluorescence emission spectroscopy acquisition.

**Figure S3.**
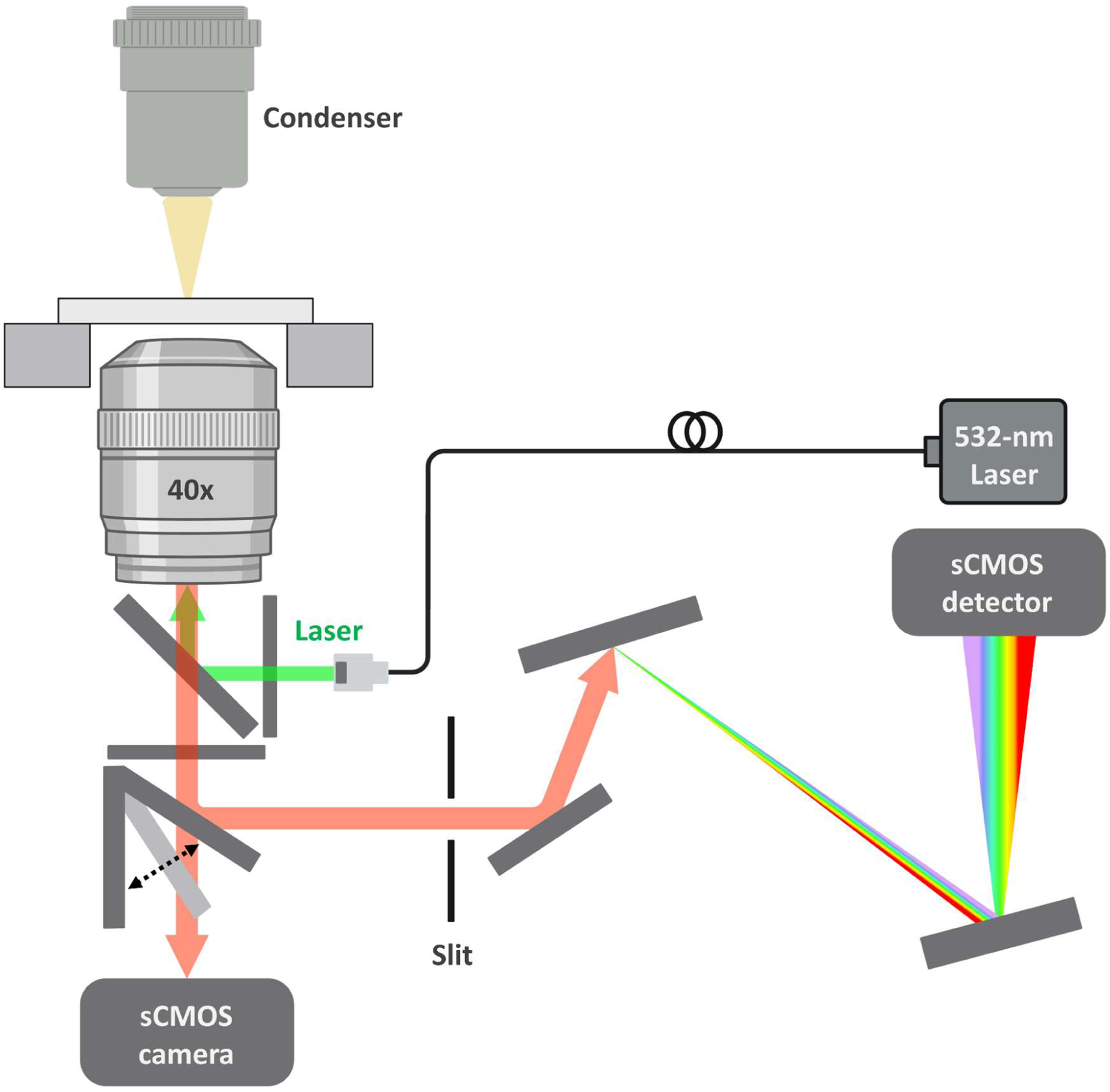
Custom-built optical system for *in situ* dark-field and fluorescence imaging with single-particle spectroscopy.

#### Supplementary Note S11: FDTD stimulation

All FDTD simulations were performed using Ansys Lumerical 2020 R2.2 (Ansys, Inc., Canada). A plane wave was employed as the incident light to calculate electric field distribution as well as excitation and decay rate enhancements, with a mesh size of 0.1 nm. Perfectly match layer (PML) boundary conditions were used for all the calculations. The ND was modeled with a refractive index of 2.42, gold was described using dielectric functions from CRC, and the surrounding medium was assigned a refractive index of 1.0.

Enclosed plasmonic ND was modeled to consist of a spherical ND (diameter: 40 nm) as the core seamlessly surrounded by a cluster of Au NSs (diameter: 90 nm). As a control, a conventional plasmonic configuration (i.e., semi-enclosed plasmonic ND) was modeled, which consists of a spherical ND (diameter: 40 nm) as the core surrounded by several Au nanospheres (diameter: 80 nm). Modeling the ND as a spherical shape is based on the consideration of balancing all the possible shapes of NDs. A single NV center was assumed in the ND and modeled as an electric dipole oscillating horizontally with an intrinsic quantum efficiency of 0.7. LSPR hotspots and the corresponding electric field enhancements were visualized at the major emission wavelength (695 nm) of NV center.

The excitation and decay rate enhancements were calculated as a function of the NV center position within the nanocavity. The simulations included both radiative and non-radiative decay channels, and the Purcell factor was extracted from the total power emitted by the dipole relative to its emission in a homogeneous medium. The position-dependent calculations account for the potential locations of multiple NV centers within the ND. Position-dependent quantum efficiency is derived from the radiative-to-total decay ratio.

**Figure S4.**
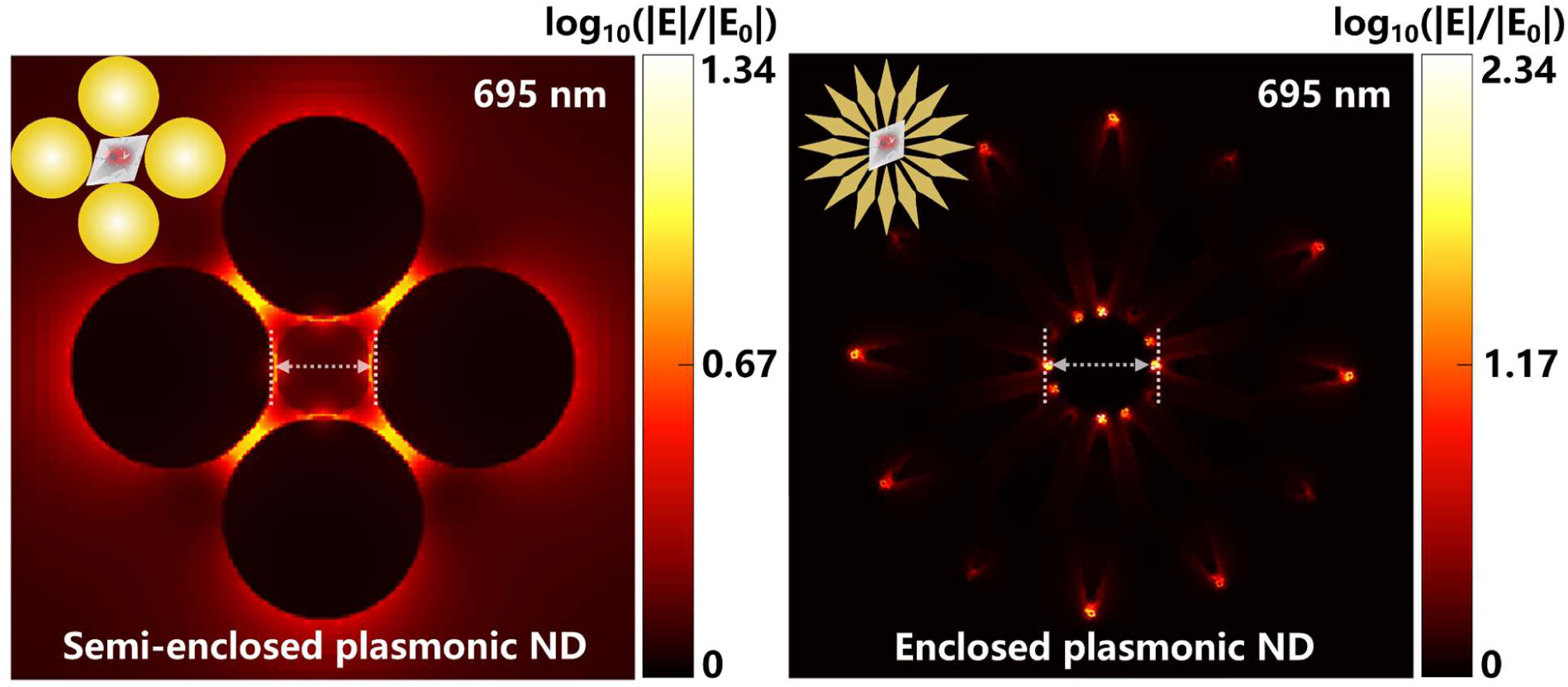
Calculated electric field enhancement for semi-enclosed plasmonic ND (control) and enclosed plasmonic ND at the major emission wavelength (695 nm) of NV center.

#### Supplementary Note S12: Fluorescence lifetime imaging microscopy (FLIM)

FLIM characterizations were conducted using a MicroTime200 system (PicoQuant, Germany) integrated with an Olympus inverted microscope. Excitation was provided by a 520 nm pulsed laser diode operating at a 5 MHz repetition rate. The excitation intensity was carefully regulated using a primary variable neutral density filter (Lbtek, NDFR-50S-3M). The laser beam sequentially traversed a 500 nm short-pass filter (Jcoptix, OFE1SP-500) and a 520 nm dichroic mirror before being focused onto the sample via a 60× oil immersion objective (NA = 1.2). The back-collected PL signal was reflected by the same dichroic mirror, spectrally isolated by a 650-720 nm band-pass filter, and routed through a secondary variable neutral density filter and a 300 μm confocal pinhole. Ultimately, the photons were detected by a single-photon avalanche diode (SPAD). To strictly avoid pile-up effects, the secondary neutral density filter was tuned to maintain the photon counting rate below 3% of the excitation rate. Time-correlated single-photon counting (TCSPC) data were acquired via a PicoQuant TimeHarp 260 module. Spatial mapping was executed with a 50 nm step size and a 50 ms dwell time per pixel. The excitation power at the objective was calibrated using a Thorlabs S171C slide-mounted sensor. PicoQuant SymphoTime 64 software was employed to generate and normalize the PL intensity and lifetime maps, with all subsequent quantitative analyses performed in OriginLab.

**Figure S5.**
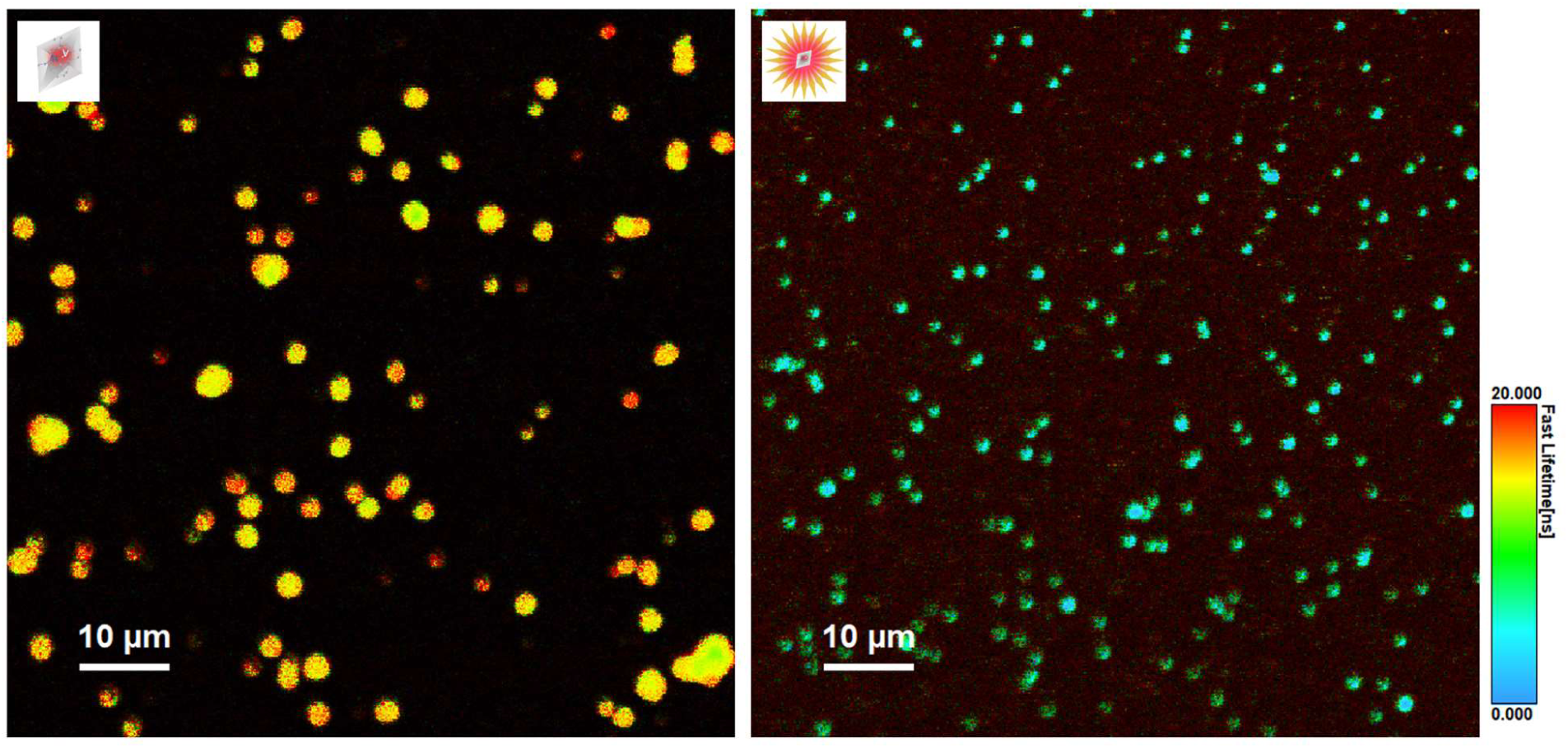
Large-area FLIM images (scale bar: 10 µm) of NDs and enclosed plasmonic NDs.

#### Supplementary Note S13: Photon emission saturation measurements

Photon emission from the sample was excited using a 532 nm laser (OBIS Laser Box, Coherent Inc., USA) with the power varied from 0 to 3 mW. The emitted photons were collected and detected by a single-photon counting module (SPCM-AQRH-14, Excelitas, USA). Time-resolved photon arrival events were recorded using a time-to-digital converter (Time Tagger, Swabian Instruments, Germany) and streamed to a computer for acquisition and analysis. Saturation curves of photon emission as a function of excitation power were generated and fitted using a logistic function to extract the saturation behavior of the emitters.

### Section 6: Wide-field optically detected magnetic resonance (ODMR) measurement

#### Supplementary Note S14: Fabrication of microwave antenna chip

The omega-shaped microwave antenna chip was fabricated via a three-step process: (1) the deposition of titanium-gold (Ti-Au) bilayer on the borosilicate glass wafer via electron beam evaporation; (2) the fabrication of omega-shaped photoresist pattern on the wafer via standard maskless lithography; and (3) the fabrication of the wafer deposited with an omega-shaped Ti-Au bilayer (i.e., microwave antenna chip) via wet etching.

Firstly, a 50 nm-thick Ti layer and a 500 nm-thick Au layer were sequentially deposited on the surface of a 500 μm-thick borosilicate glass wafer (BF33, Schott AG, Germany) by electron beam evaporation via an E-beam evaporation system (Vapour station 4, Oxford Vacuum Science, UK). Ti layer served as an adhesive layer to ensure the strong adhesion between Au layer and the wafer. Post-deposition, the wafer was allowed to cool down under vacuum to avoid oxidation of the freshly deposited layers. Subsequently, the entire wafer was segmented into smaller wafers of the desirable size via a wafer scribing and cleaving system (LatticeAx 420, Ted Pella, USA). These wafers were then immersed in acetone for 10 min, followed by anhydrous ethanol for 5 min, and finally rinsed with ultrapure water.

To fabricate the omega-shaped pattern, the positive photoresist AZ 5214 was spin-coated onto the wafer deposited with Ti-Au bilayer at 600 rpm for 5 s and another 4000 rpm for 30 s to deposit a 1.3 µm-thick photoresist coating. The wafer was then baked in 100 °C for 3 min. Subsequently, UV exposure (300 mJ cm^-2^) was applied to selectively irradiate the photoresist coating on the wafer, excluding the regions corresponding to the desired pattern, via a maskless aligner (MLA150, Heidelberg Instruments, Germany). Afterwards, the wafer was developed in AZ 300MIF for 37 s under continuous shaking to dissolve the exposed photoresist. The residual photoresist was washed away via ultrapure water and dried with nitrogen gas. The wafer was then heated at 100 °C for 90 s to enhance the stability of the photoresist pattern.

The wafer, deposited with Ti-Au bilayer and photoresist pattern, was immersed in a solution containing iodine (60 mg mL^-1^) and potassium iodide (200 mg mL^-1^) for 20 s to selectively etch the Au layer beyond the photoresist pattern. This was followed by rinsing with ultrapure water and drying with nitrogen gas. Subsequently, the wafer was immersed in 36% concentrated hydrochloric acid for 40 min to selectively etch the Ti layer beyond the photoresist pattern, followed by rinsing with ultrapure water and drying with nitrogen gas. Afterwards, the wafer was immersed in N-methylpyrrolidone at 60 °C for 30 min and then in acetone for an additional 30 min to remove the photoresist pattern. Finally, the wafer was rinsed with isopropanol and ultrapure water, and dried with nitrogen gas, resulting in the fabrication of a wafer deposited with an omega-shaped Ti-Au bilayer (i.e., microwave antenna chip).

**Figure S6.**
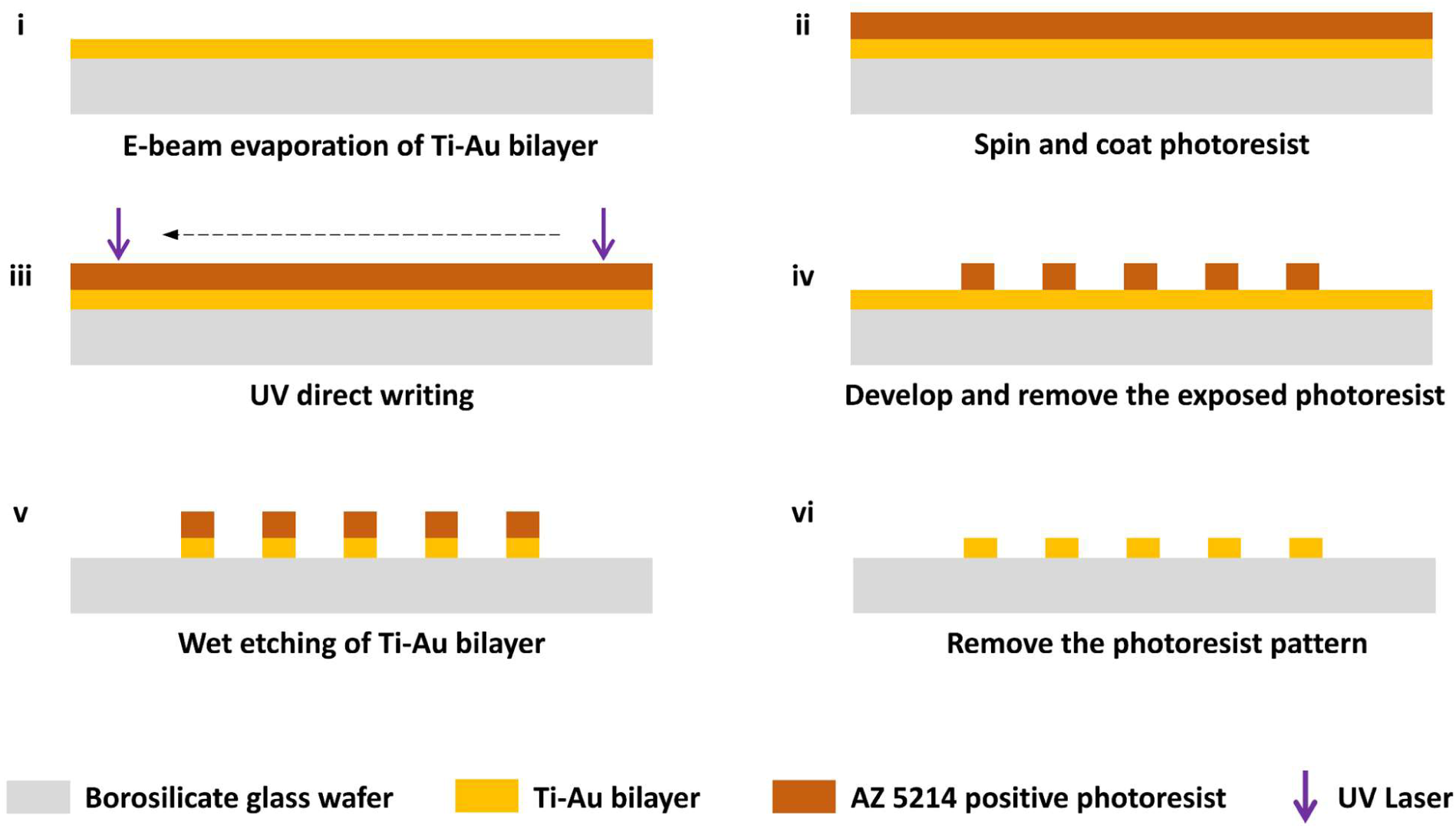
Flow chart for fabricating a microwave antenna chip.

**Figure S7.**
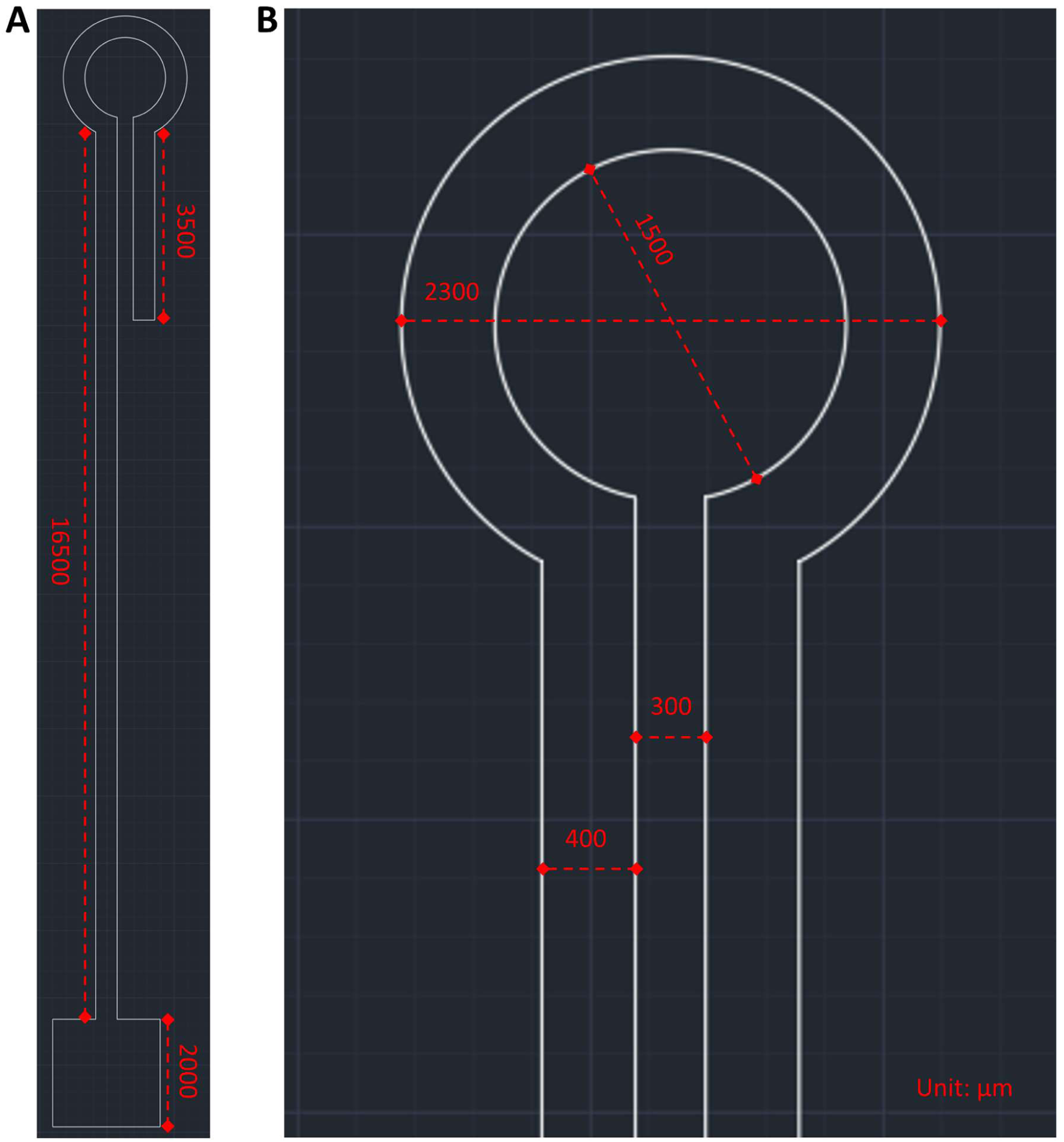
Structural parameters of the omega-shaped microwave antenna. **(A)** Overall structural parameters of the microwave antenna, including the main body and connection interface. **(B)** Magnified view of the omega-shaped region with detailed structural parameters. Unit: µm.

**Figure S8.**
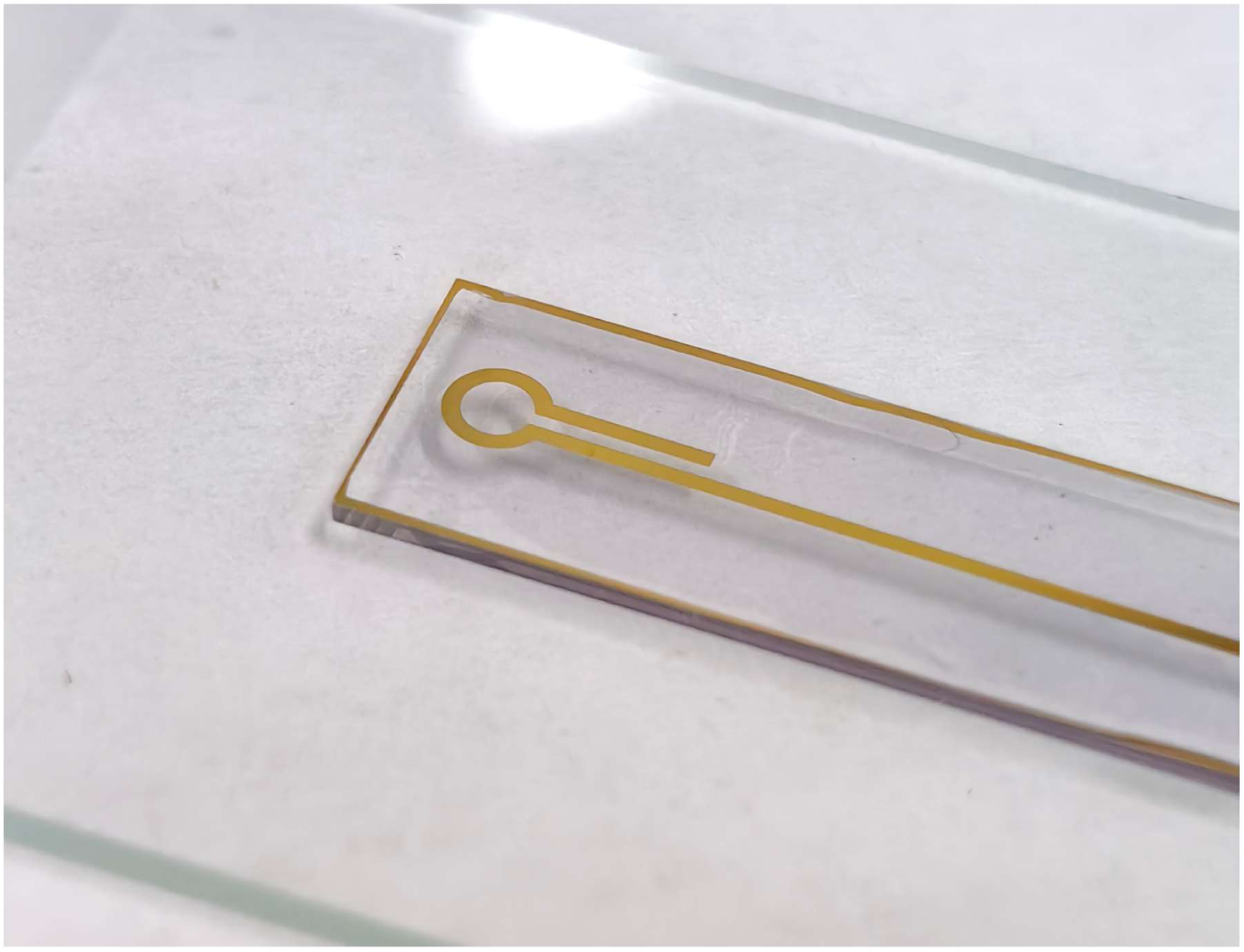
The developed microwave antenna chip.

#### Supplementary Note S15: Construction of microwave generating device

The microwave generating device is constructed from a RF signal source, a power amplifier, a DC power supply, a RF isolator, and a microwave antenna chip. RF signal source (SSG3032X-IQE, Siglent Technologies, China) is employed to produce RF signal sequences at preset frequencies. Power amplifier (ZHL-16W-43-S+, Mini-Circuits, USA) serves to enhance the power of RF signals with a gain of 45 dB, energized by a DC power supply (GPS305D, Wanptek, China). RF isolator (UIYBCI4040A, UIY Inc., China) is integrated to prevent the reflection of electrical signals to protect critical components. Microwave antenna chip is used for emitting microwaves at preset frequencies and facilitating their focused and uniform propagation.

**Figure S9.**
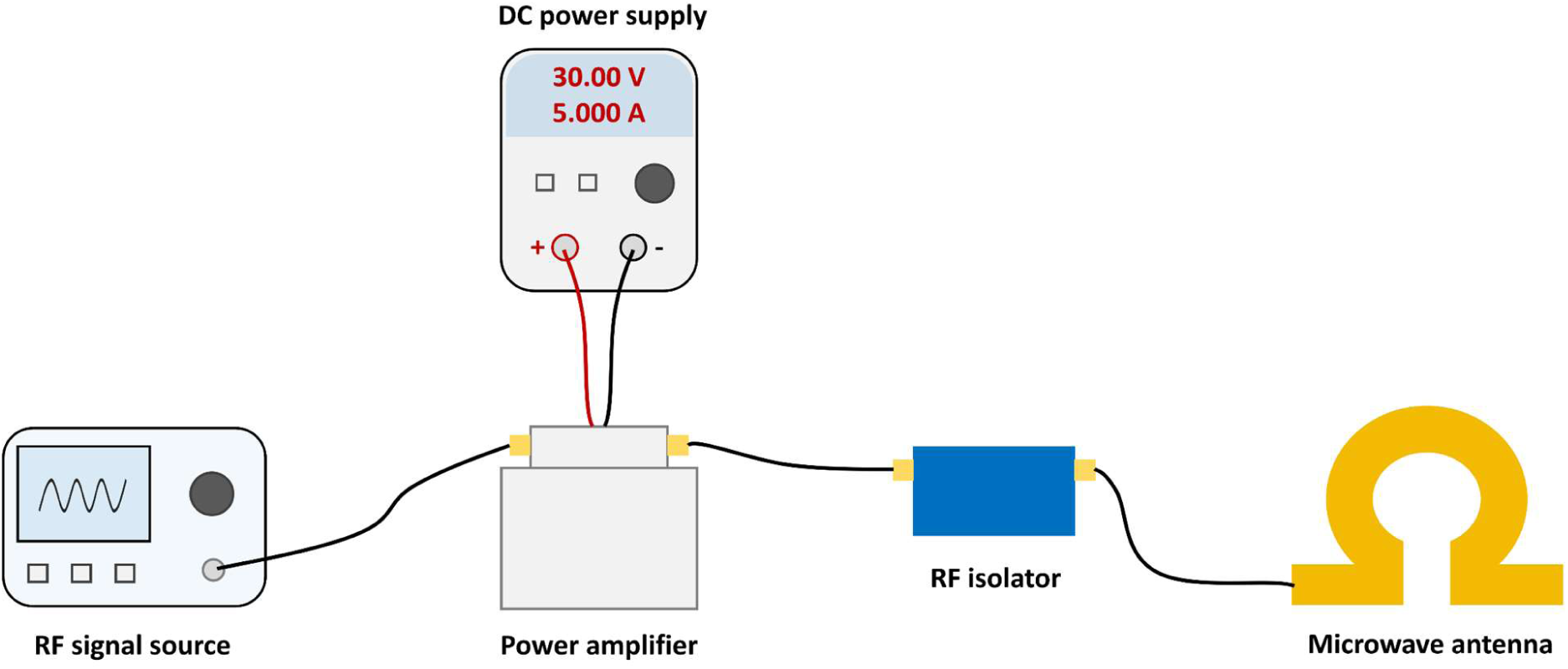
Schematic of the microwave generating device.

#### Supplementary Note S16: Wide-field ODMR measurement protocol

Wide-field ODMR measurements were performed on an inverted fluorescence microscope configured in total internal reflection fluorescence (TIRF) geometry to selectively excite NV centers located near the substrate interface. A continuous-wave 532 nm laser was used for optical excitation, and the emitted fluorescence was collected through a high-numerical-aperture objective and recorded using a scientific CMOS camera. The excitation power at the sample plane was maintained constant for all measurements to ensure comparable spin readout conditions.

Microwave irradiation was delivered through a lithographically patterned microwave antenna positioned adjacent to the sensing substrate, while a movable permanent magnet provided a tunable static magnetic field. During measurements, microwave power was fixed at 50 mW unless otherwise specified. The microwave frequency was scanned from 2820 MHz to 2920 MHz in 1 MHz steps with a dwell time of 50 ms per point, and fluorescence images were synchronously acquired at each frequency to construct ODMR spectra.

The underlying spin physics follows the established NV center ground-state structure, in which resonance microwave excitation drives transitions between the ground-state spin sublevels *m_s_*=0 and *m_s_*=±1. In the presence of an external magnetic field, the degeneracy of *m_s_*=±1 states is lifted via the Zeeman effect, giving rise to two symmetrically split resonance frequencies corresponding to the *m_s_*=+1 and *m_s_*=-1 transitions. During the optical excitation and decay, the *m_s_*=±1 excited-state levels exhibit a higher propensity to undergo non-radiative decay into a metastable ‘dark’ state, leading to reduced fluorescence emission and thereby enabling optical readout of spin resonance.

ODMR spectra were obtained by averaging the fluorescence intensity over regions of interest corresponding to individual ND emitters or ensembles of emitters. The normalized fluorescence contrast was calculated as (*I*_off_ − *I*)/*I*_off_, where *I* represents the fluorescence intensity at each microwave frequency and *I*_off_ is the off-resonance fluorescence baseline. To quantify spectral parameters, ODMR resonance dips were fitted using Lorentzian functions to extract the resonance frequency, full width at half maximum (FWHM), and ODMR contrast. For measurements performed under applied magnetic fields, the separation between the two Lorentzian minima was used to determine the Zeeman splitting of the *m_s_* = ±1 transitions. All measurements were performed under identical optical excitation, microwave power, and magnetic-field alignment conditions to enable direct comparison between NDs and enclosed plasmonic NDs.

#### Supplementary Note S17: Magnetic sensitivity calculation

The magnetic sensitivity (5) can be calculated by the following equation:

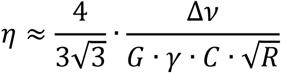

Where 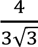 refers to the numerical parameter related to the Lorentzian profile of spin resonance, Δν is the FWHM of ODMR spectrum, *G* is the gain factor related to NV centers (≈ 2), γ is the electron gyromagnetic ratio (= 28 MHz/mT), *C* is the ODMR contrast, and *R* is the saturated photon emission rate.

**Figure S10.**
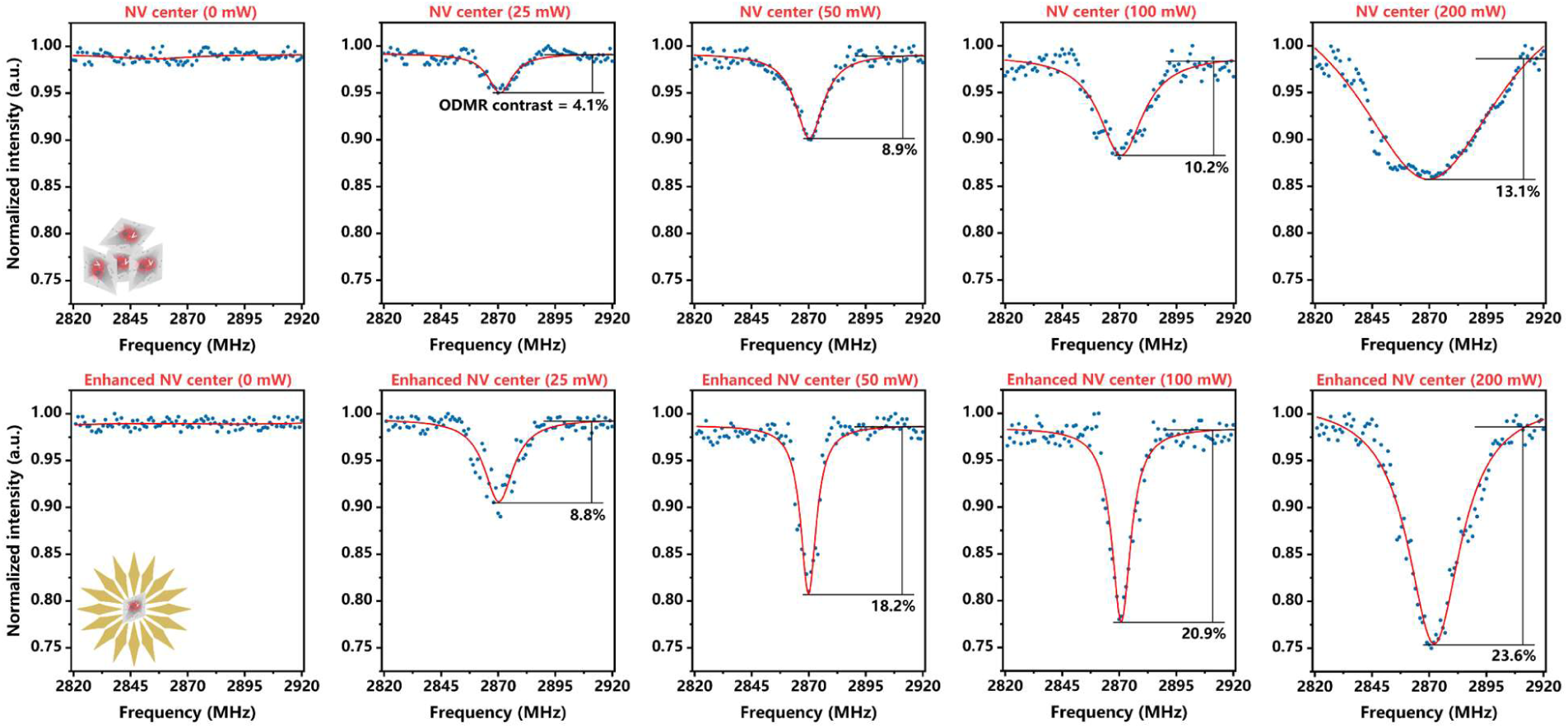
ODMR measurements of NV centers and enhanced NV centers at increasing microwave power (0-200 mW) and zero magnetic field. Enhanced NV centers exhibit consistently higher ODMR contrast and narrower linewidths than NV centers.

**Figure S11.**
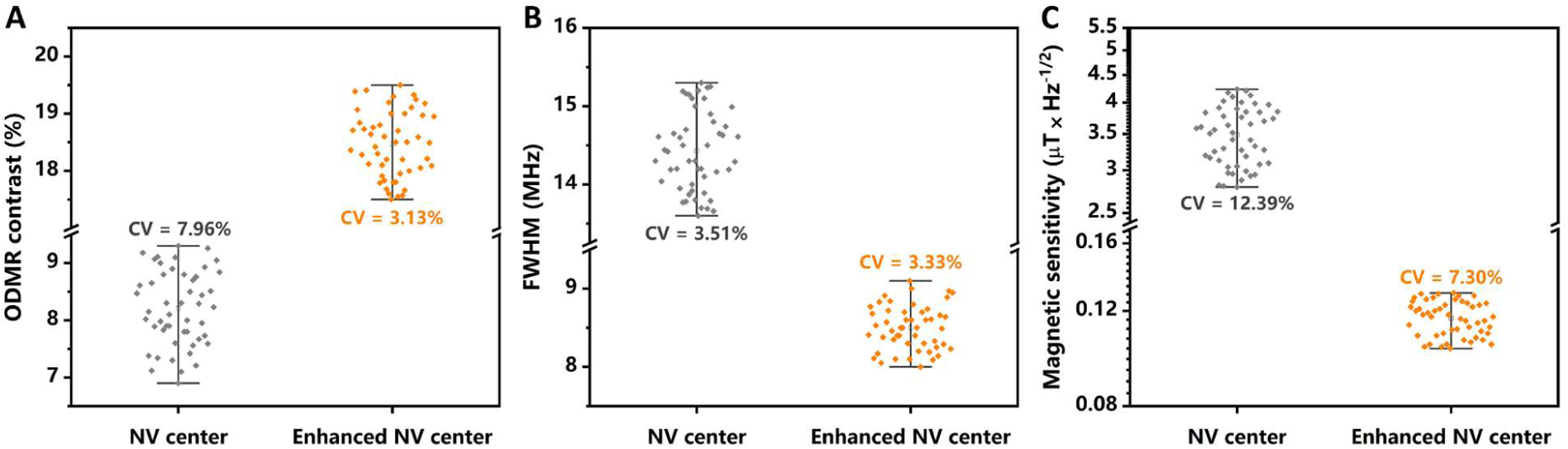
Spin-state readout performance statistics of NV centers versus enhanced NV centers: **(A)** ODMR contrast, **(B)** FWHM, and **(C)** derived magnetic sensitivity.

### Section 7: Digital event statistics and BISPIN quantification validation

#### Supplementary Note S18: Definition of the BISPIN decision threshold

The digital classification threshold used in the BISPIN framework was determined from blank-control measurements to statistically separate noise-induced splitting fluctuations from signal-associated events. Specifically, ODMR spectra were first acquired under control conditions in which no external magnetic perturbation or bio-spin activity was present. For each sensing location, the resonance splitting (Δ) was extracted using the same ODMR fitting pipeline described above, generating a baseline distribution of splitting values arising from instrumental noise, fitting uncertainty, and intrinsic NV variability.

The decision threshold (Δ_th_) was then defined using a five-sigma statistical criterion applied to this baseline distribution:

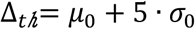

where *μ*_0_ and *σ*_0_ denote the mean and standard deviation of the splitting values measured in the blank control dataset. This threshold represents the upper bound of the baseline fluctuation range with a confidence level corresponding to the 5σ statistical criterion.

In subsequent measurements, each ODMR-derived splitting value (Δ) was compared to the predefined threshold Δ_th_. Events satisfying Δ ≥ Δ_th_ were classified as positive events, indicating statistically significant splitting excursions beyond baseline noise, whereas events with Δ < Δ_th_ were classified as negative events. This threshold-based binarization converts analog ODMR readouts into digital event outcomes, enabling statistical analysis of stochastic magnetic fluctuations through event-counting metrics.

**Figure S12.**
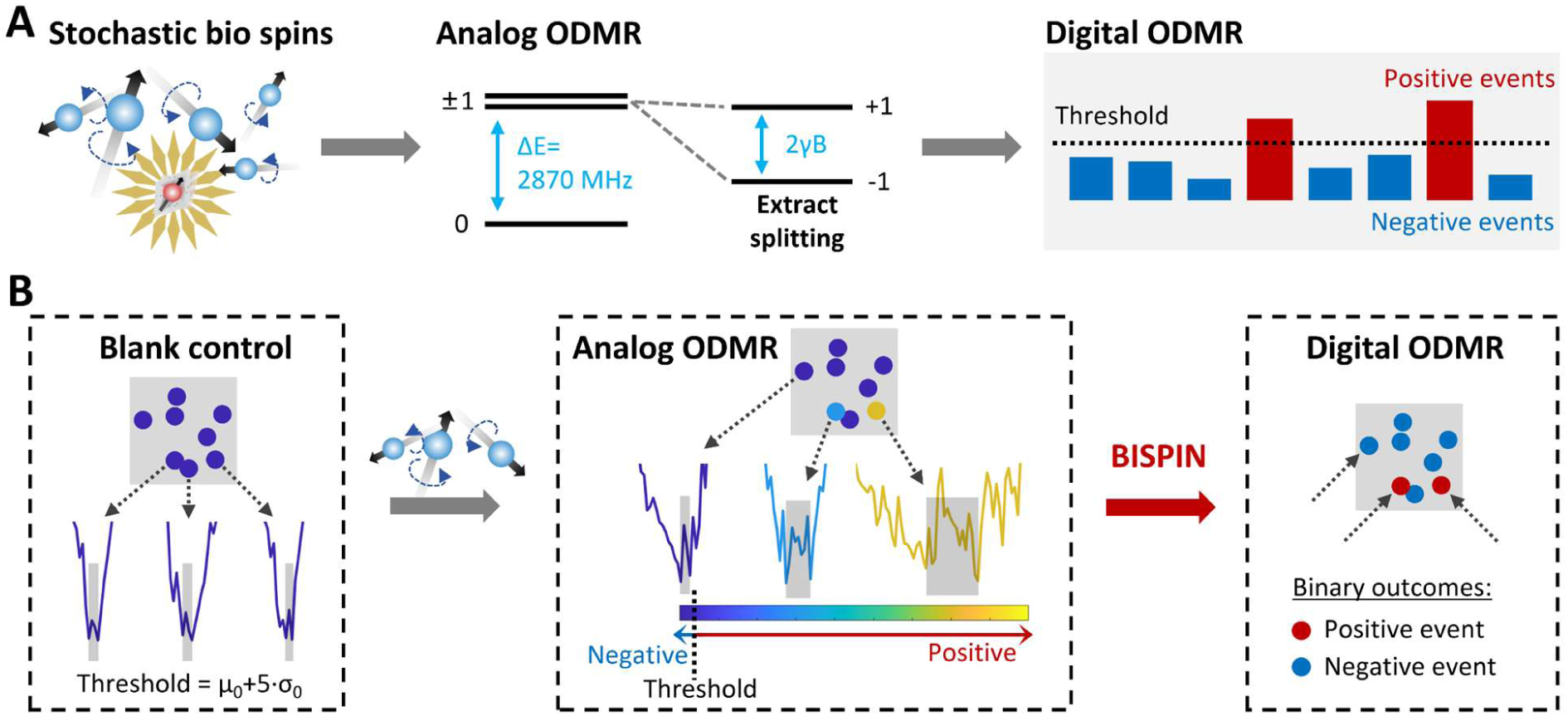
BISPIN workflow. **(A)** Stochastic bio-spin fluctuations are recorded via analog ODMR, from which resonance splittings are extracted. These analog signals are then converted into digital positive and negative events by applying a predefined threshold. Specifically, **(B)** ODMR spectra are first acquired under blank control, defining the threshold Δ_t_*_ℎ_*= *μ*_0_ + 5 · *σ*_0_. Under stochastic bio-spin conditions, each ODMR-derived splitting value (Δ) is compared to the threshold Δ_th_. Events satisfying Δ ≥ Δ_th_ are classified as positive events, and as negative events otherwise. This threshold-based binarization converts analog ODMR readouts into digital event outcomes, enabling statistical analysis of stochastic magnetic fluctuations via event-counting metrics.

#### Supplementary Note S19: Validation of digital threshold statistics

Controlled static magnetic fields were applied to NV and plasmon-enhanced NV substrates using a calibrated, movable permanent magnet. Field strengths of 0.1, 0.3, and 0.5 mT were sequentially applied, and verified with a Hall probe prior to each acquisition. Wide-field ODMR imaging was performed across multiple fields of view at each applied field to acquire statistically significant datasets.

For each sensing location, resonance splittings (Δ) were extracted using the established ODMR fitting pipeline. Positive-event counts were determined by comparing each Δ to the predefined BISPIN decision threshold (Δ_th_) established from blank-control measurements. The positive-event fraction, p = P(Δ ≥ Δ_th_), was calculated for each field strength and substrate type.

Absolute event counts and relative standard deviations (RSDs) were analyzed to assess both the mean event response and the statistical fidelity of the digital observable. Reduced RSDs indicate convergence toward Poisson-like counting statistics, whereas elevated variability reflects sparse positive events or increased sensitivity to fitting uncertainty. Comparisons between NV and plasmon-enhanced NV substrates quantified the improvement in event detection reliability and convergence of BISPIN statistics under weak-field conditions relevant for cellular measurements.

#### Supplementary Note S20: Set-up of BISPIN microscopy

Enhanced NV array substrate is mounted on the stage of BISPIN microscope integrating a wide-field fluorescence microscope with a microwave generating device to reconstruct stochastic magnetic perturbations induced by bio spins.

Wide-field ODMR imaging is performed on an inverted fluorescence microscope configured in TIRF geometry to selectively excite the single NV layer. A continuous-wave 532 nm laser is used for optical excitation, filtered through a 532±2 nm band-pass filter and reflected by a 532 nm long-pass dichroic mirror. The beam is focused onto the sample through a 100× oil immersion objective (NA=1.49). The emitted fluorescence is collected by the same objective, separated from the excitation laser by the dichroic mirror, passed through a 610 nm long-pass filter to suppress residual scattering, and recorded by a back-illuminated sCMOS camera.

Microwave irradiation is delivered through a lithographically patterned microwave antenna positioned adjacent to the sensing substrate, driven by the microwave generating device consisting of RF signal source, power amplifier, and RF isolator. During measurements, microwave power is fixed at 50 mW unless otherwise specified. The microwave frequency is scanned from 2820 MHz to 2920 MHz in 1 MHz steps with a dwell time of 50 ms per point, and fluorescence images are synchronously acquired at each frequency to construct ODMR spectra, followed by BISPIN analysis.

**Figure S13.**
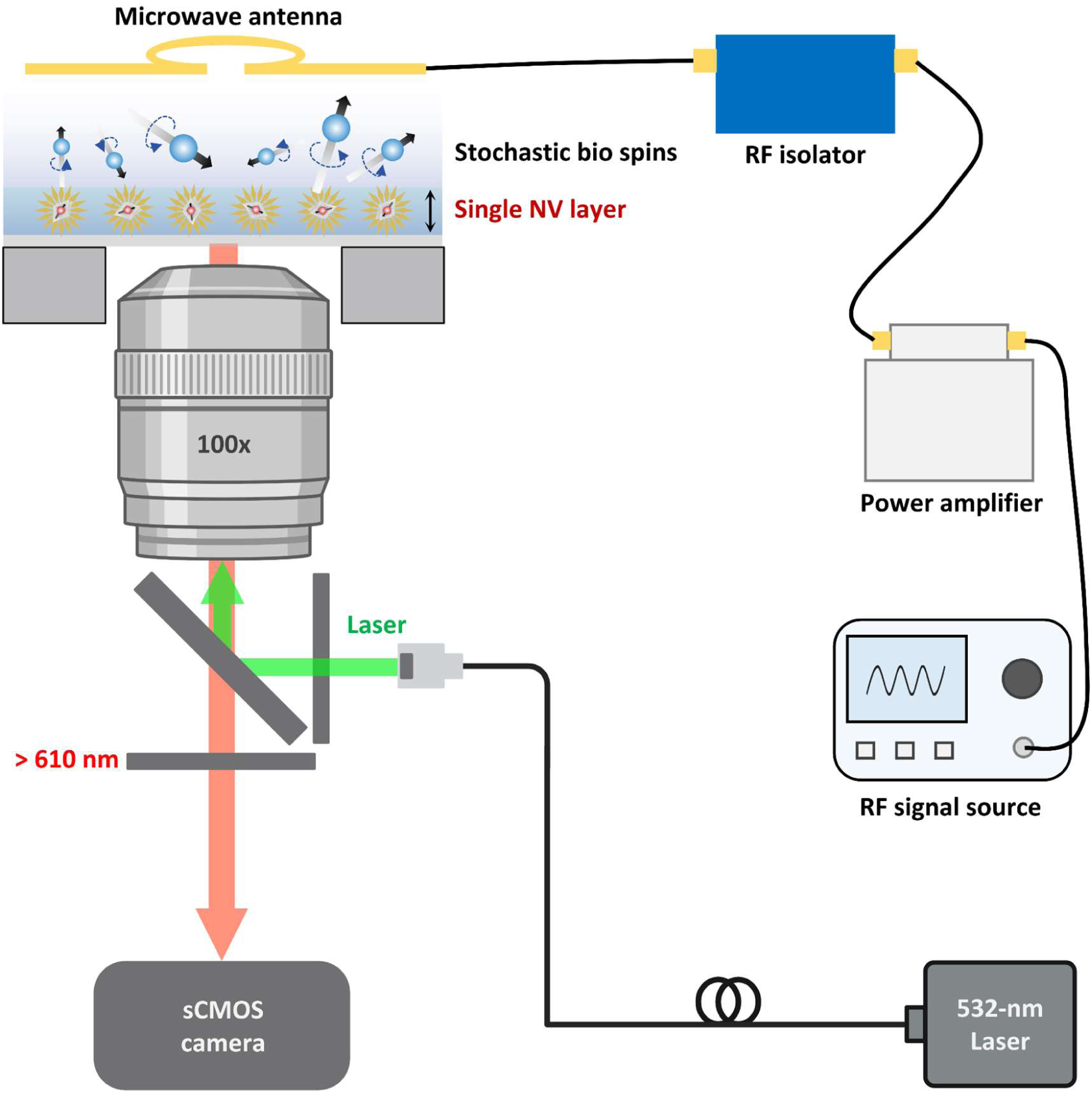
Set-up of BISPIN microscopy. Enhanced NV array substrate is mounted on the stage of BISPIN microscope integrating a wide-field fluorescence microscope with a microwave generating device for reconstruction of stochastic magnetic perturbations induced by bio-spins. TIRF-mode ODMR imaging is performed to selectively excite a single NV layer. Fluorescence image of substrate is recorded by a back-illuminated sCMOS camera. Microwave irradiation is delivered via a microwave antenna adjacent to the substrate, driven by RF signal source, power amplifier, and RF isolator. ODMR spectra are acquired by sweeping the microwave frequency from 2820 to 2920 MHz with synchronized fluorescence imaging, followed by BISPIN analysis.

#### Supplementary Note S21: Spatial resolution of enhanced NV array substrate

The spatial distribution of enhanced NV array substrates was quantified to assess the effective sensor density. Wide-field fluorescence images were acquired using a 100× oil-immersion objective (NA = 1.49) on an inverted microscope. Individual NV fluorescence spots were identified and segmented using the FIJI ROI Manager. The centroids of all detected spots were computed to represent the spatial coordinates of each emitter.

For each NV fluorescence spot, the nearest-neighbor center-to-center distances were calculated, and the resulting distribution was used to evaluate the average center-to-center spacing across the substrate. This metric provides a quantitative measure for the effective sensor density and spatial sampling resolution of the platform.

**Figure S14.**
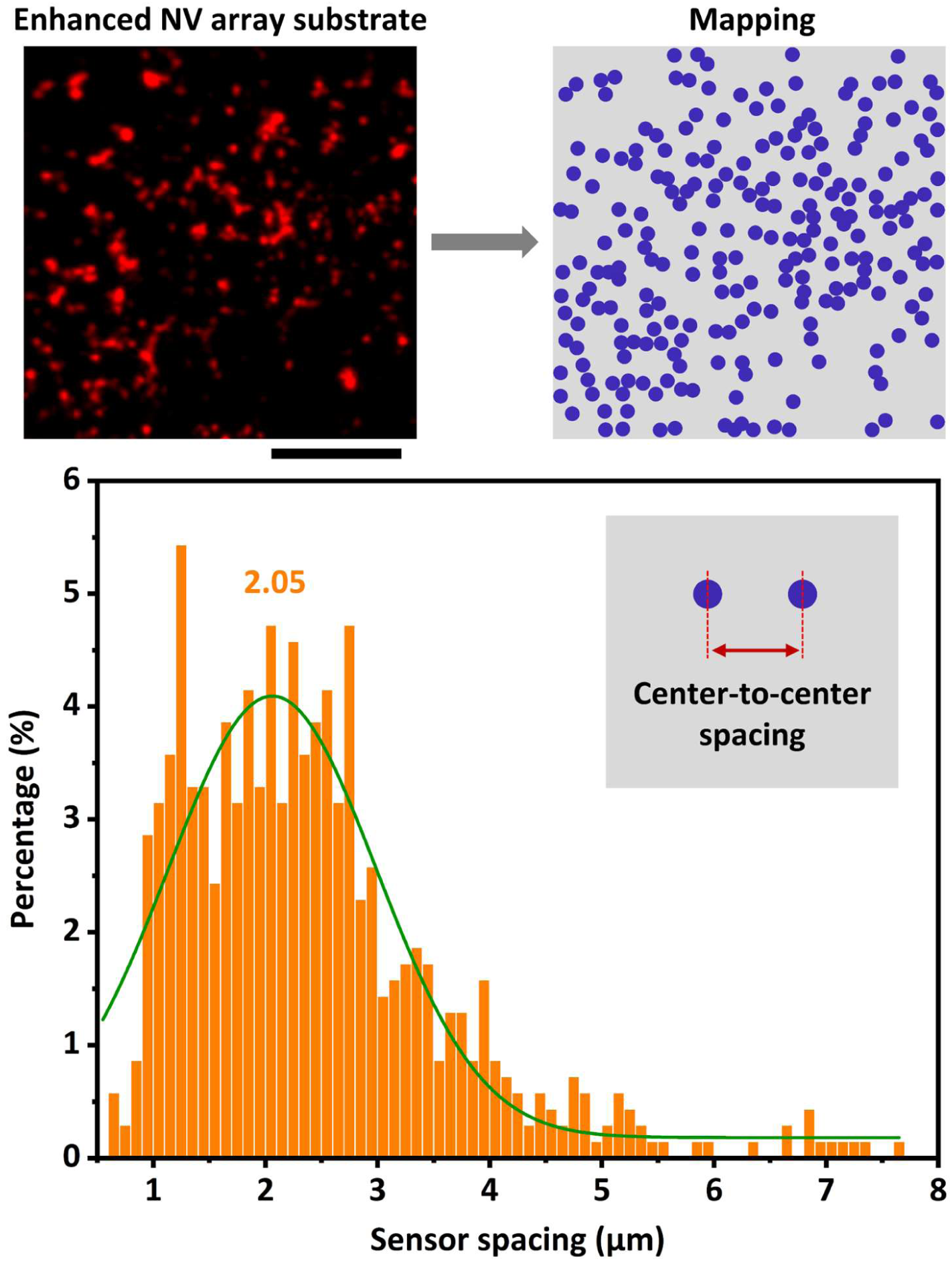
Spatial resolution of the enhanced NV array substrate. Wide-field fluorescence imaging (scale bar: 10 µm) is converted into NV emitter mapping, identifying spatial positions of all NV fluorescence spots. Nearest-neighbor center-to-center distances are calculated for each spot, and the resulting distributions (n = 700) quantify the average spacing (∼2.05 µm) between NV fluorescence spots, reflecting the effective sensor density and spatial sampling resolution of the substrate. Curve represents Gaussian fitting.

#### Supplementary Note S22: Extracellular calibration based on ion-driven magnetic noise

Enhanced NV array substrates were immersed in DMEM buffer and positioned between orthogonal electrode pairs to establish a fully controlled extracellular calibration. Two alternating electrical signals of equal input power (−20 dBm to 0 dBm) but incommensurate frequencies (9 kHz and 11.27 kHz) were applied to drive temporally varying ionic currents, generating stochastic magnetic perturbations with fluctuating magnitude and direction within the imaging area.

Wide-field fluorescence imaging and ODMR measurements were performed over a 30 μm^2^ area, approximating a single-cell-sized region. In the absence of electrode input, baseline ODMR splitting maps were acquired to verify low-noise conditions. During electrode-driven experiments, temporally continuous ODMR imaging captured the stochastic magnetic fluctuations induced by the ion currents.

For each pixel and time frame, resonance splittings (Δ) were extracted using the established ODMR fitting pipeline. Digital event maps were generated by applying the predefined BISPIN decision threshold (Δ_th_) to convert analog splitting measurements into positive and negative events. Positive-event fractions were computed for the imaging region at each input power.

Event statistics—including mean positive-event fraction, coefficient of variation (CV), and linearity with input power—were evaluated to quantify the reproducibility and statistical convergence of BISPIN readouts under stochastic, directionally mixed magnetic disturbances. This approach establishes a robust calibration framework for assessing BISPIN performance in quantifying weak and fluctuating magnetic signals at the single-cell spatial scale.

**Figure S15.**
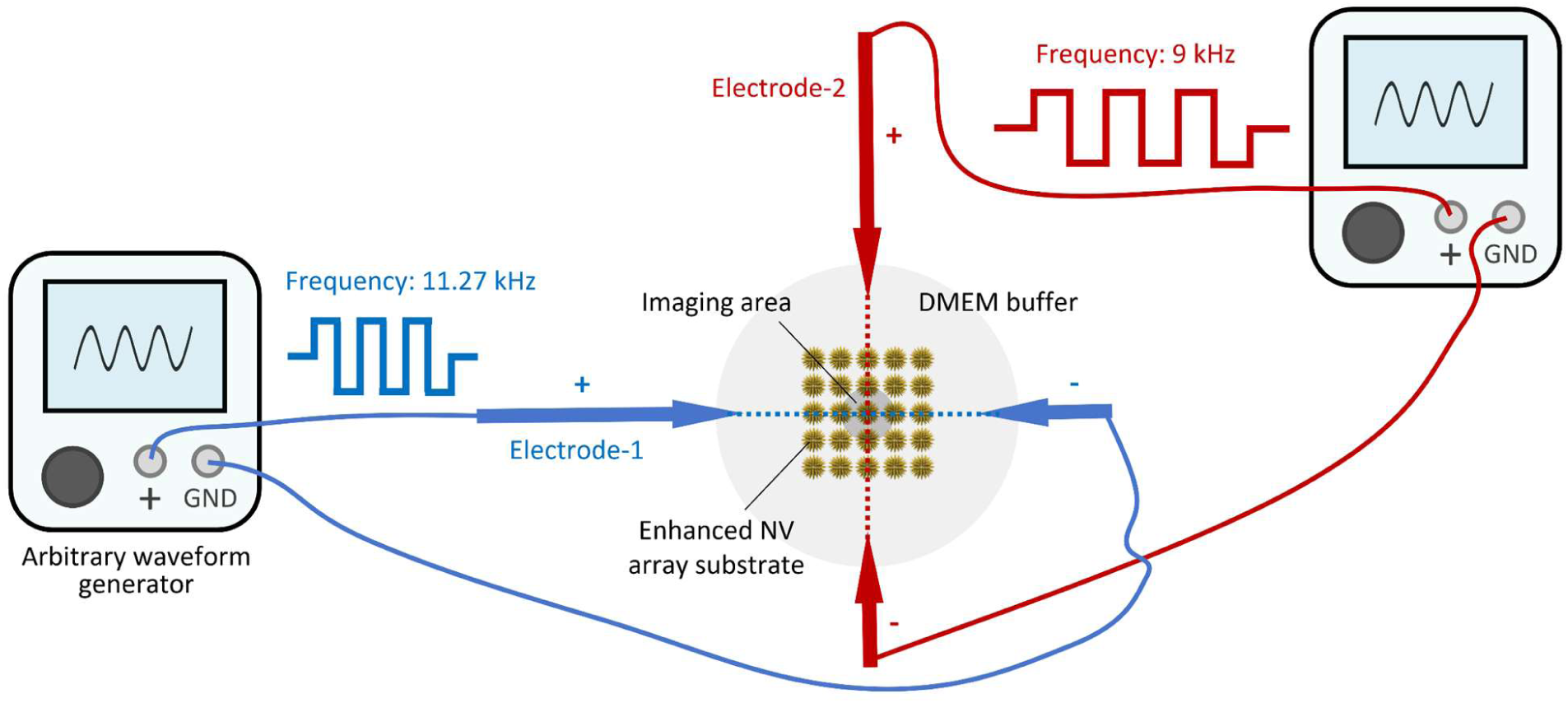
Set-up of controlled extracellular calibration. Enhanced NV array substrate is immersed in DMEM buffer and positioned between orthogonal electrode pairs. Two alternating electrical signals of equal input power (−20 dBm to 0 dBm) but incommensurate frequencies (9 kHz and 11.27 kHz) are applied to drive temporally varying ionic currents, generating stochastic magnetic perturbations with fluctuating magnitude and direction within the imaging area.

**Figure S16.**
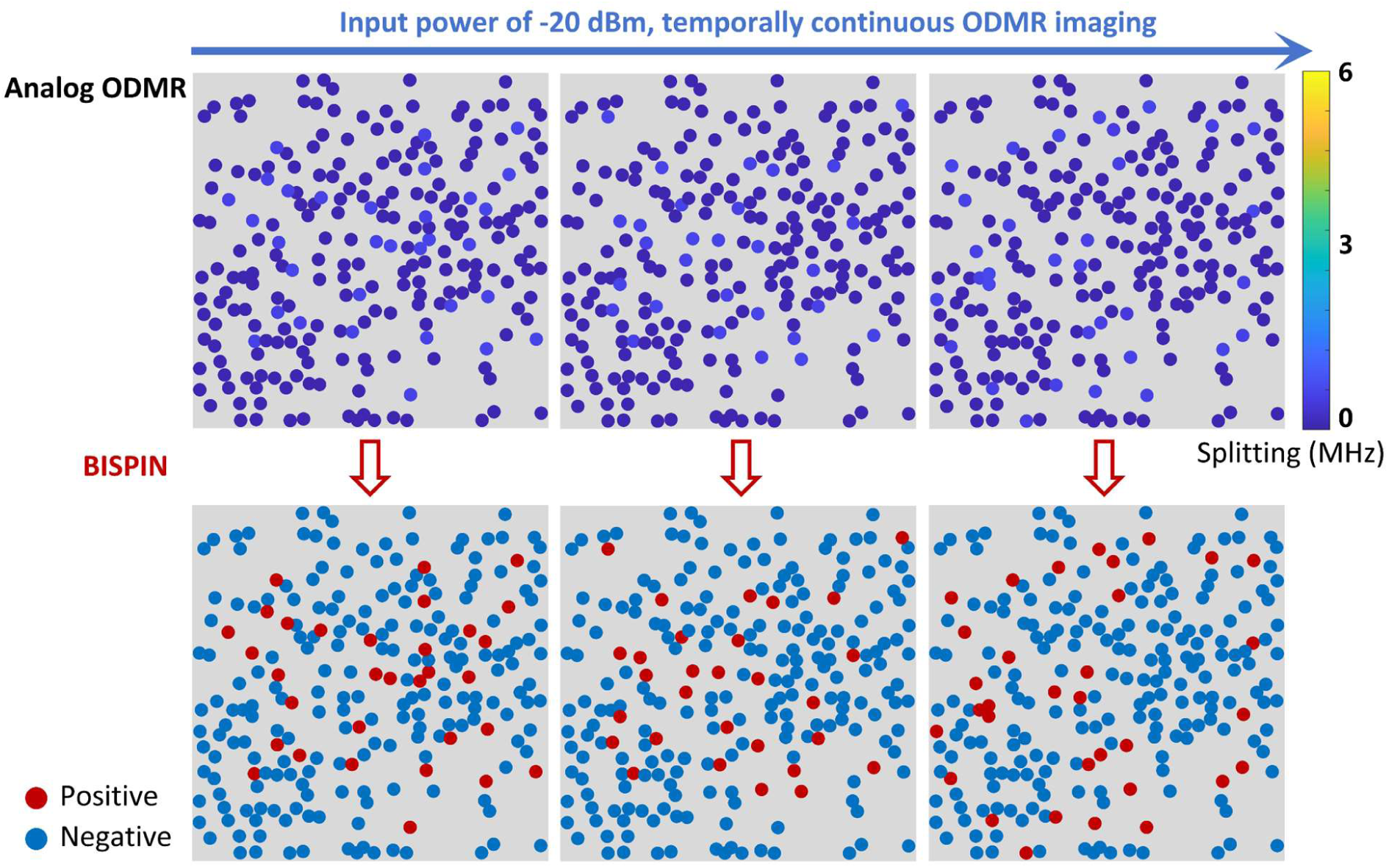
Analog ODMR splitting maps acquired under temporally continuous extracellular electrical excitation (−20 dBm input power). BISPIN microscopy converts these unstable analog readouts into binary outcomes at each pixel and time window, yielding binary event maps of positive and negative detections.

**Figure S17.**
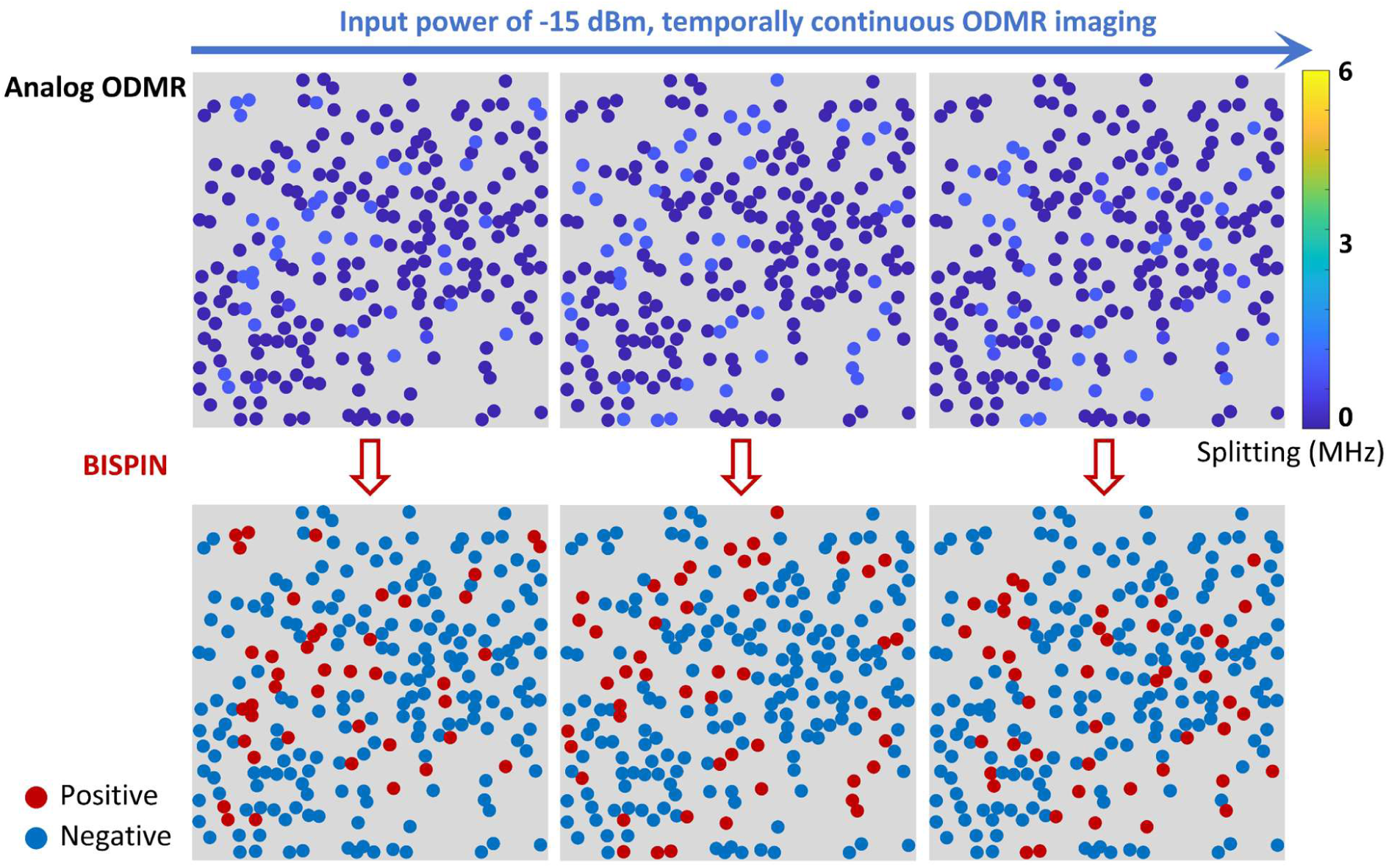
Analog ODMR splitting maps acquired under temporally continuous extracellular electrical excitation (−15 dBm input power). BISPIN microscopy converts these unstable analog readouts into binary outcomes at each pixel and time window, yielding binary event maps of positive and negative detections.

**Figure S18.**
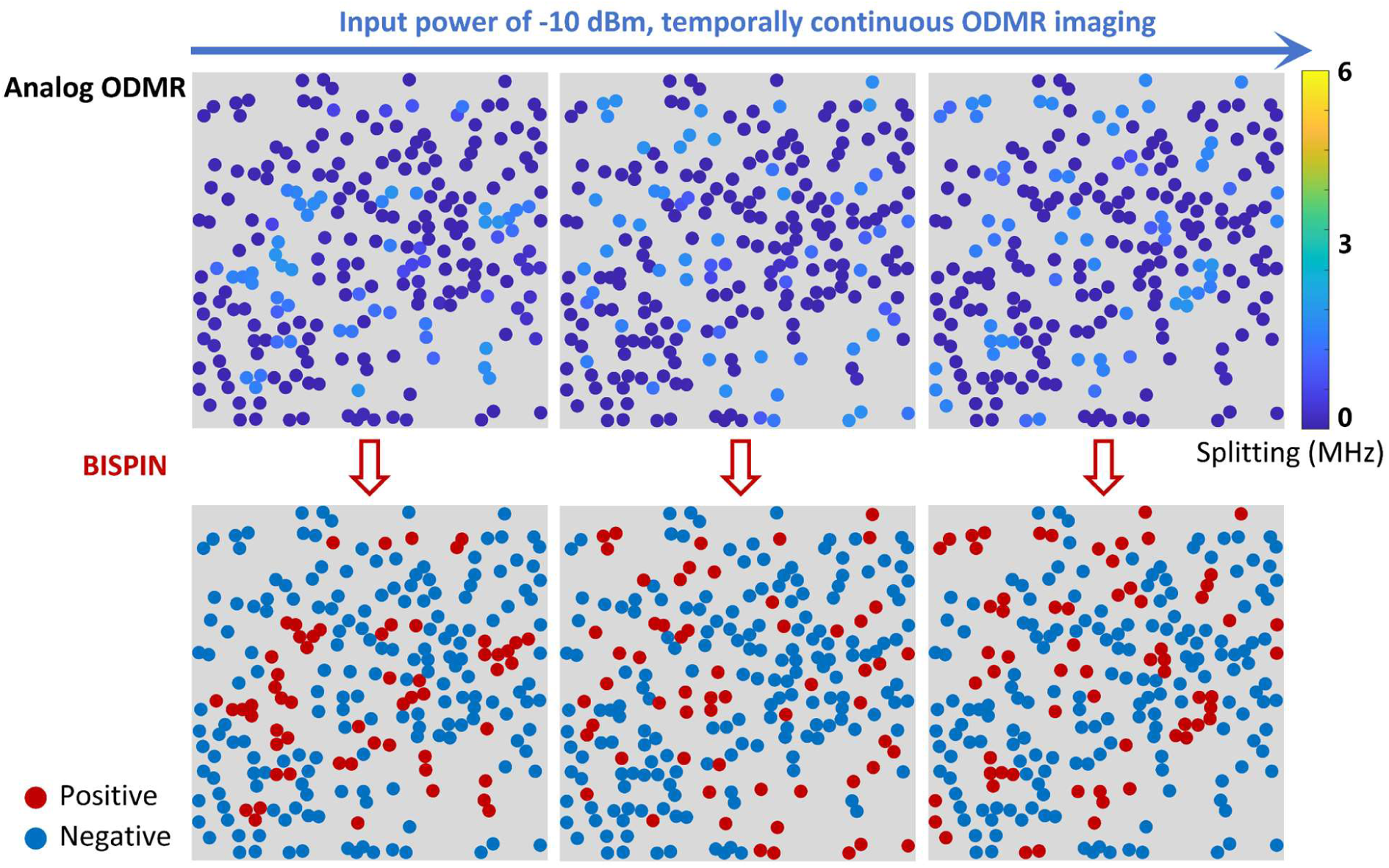
Analog ODMR splitting maps acquired under temporally continuous extracellular electrical excitation (−10 dBm input power). BISPIN microscopy converts these unstable analog readouts into binary outcomes at each pixel and time window, yielding binary event maps of positive and negative detections.

**Figure S19.**
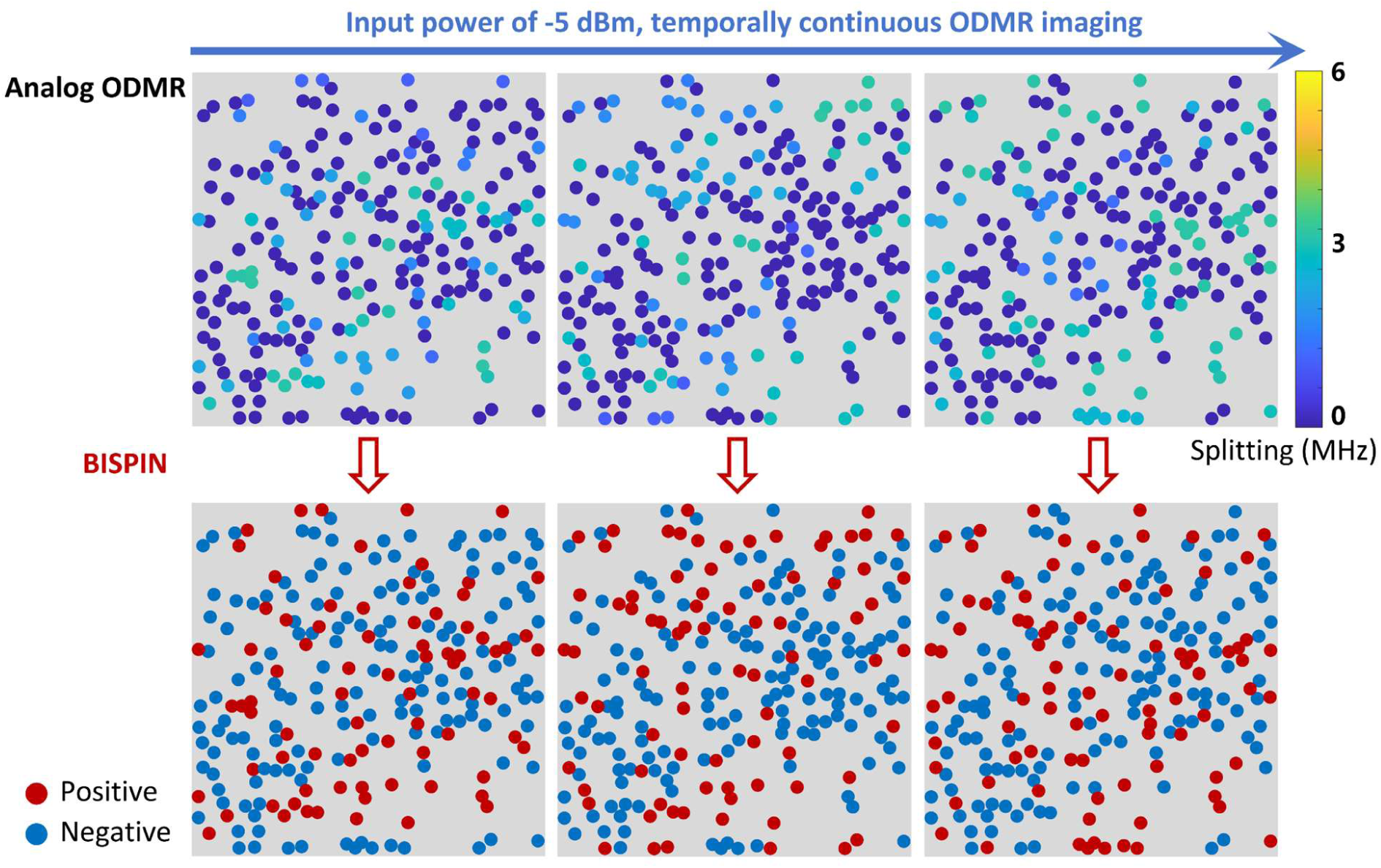
Analog ODMR splitting maps acquired under temporally continuous extracellular electrical excitation (−5 dBm input power). BISPIN microscopy converts these unstable analog readouts into binary outcomes at each pixel and time window, yielding binary event maps of positive and negative detections.

**Figure S20.**
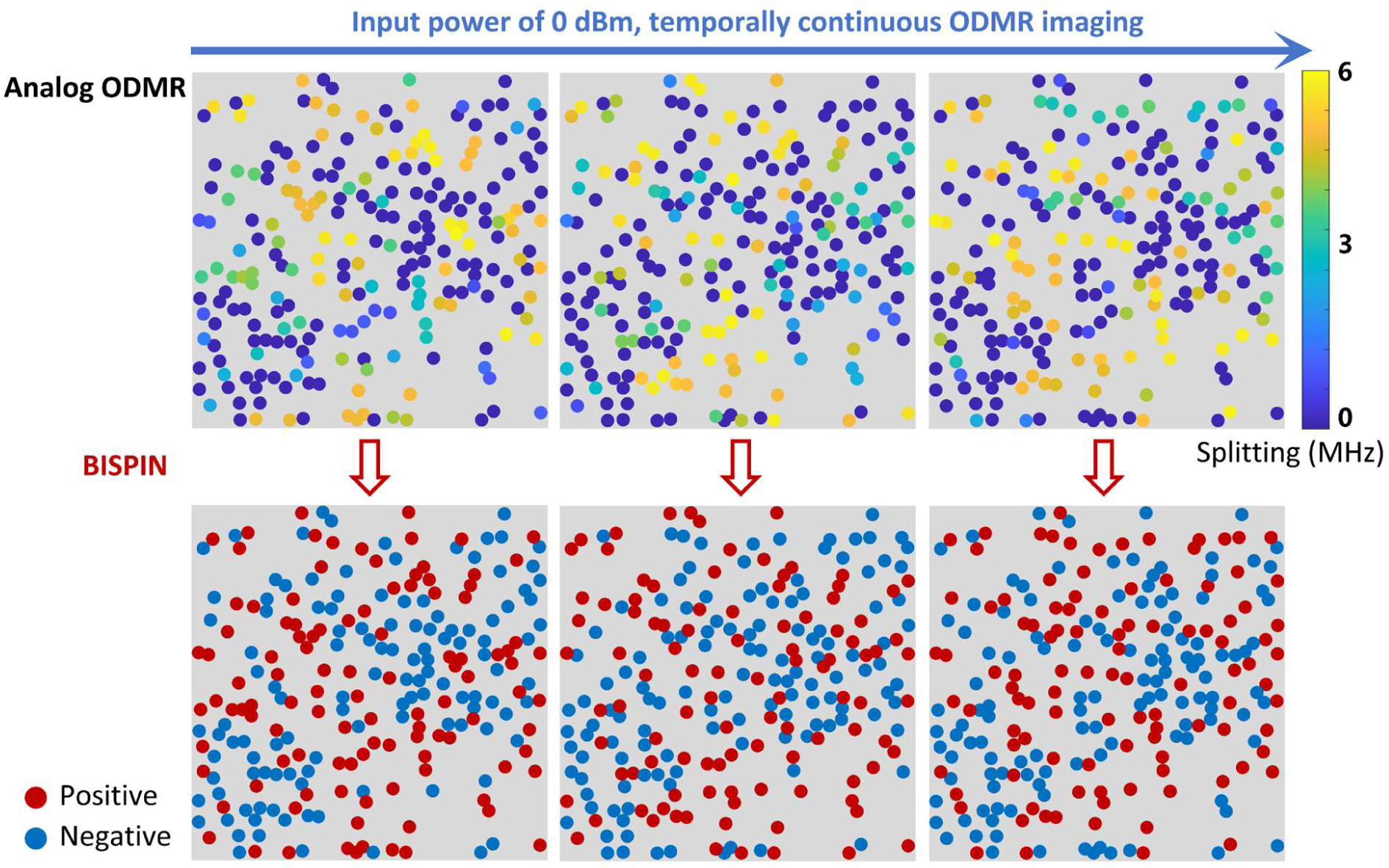
Analog ODMR splitting maps acquired under temporally continuous extracellular electrical excitation (0 dBm input power). BISPIN microscopy converts these unstable analog readouts into binary outcomes at each pixel and time window, yielding binary event maps of positive and negative detections.

### Section 8: Establishment and validation of the store-operated calcium entry (SOCE)

All solutions used in the experiments were pre-equilibrated to 37 °C by water-bath incubation for 10 min prior to use. All incubation steps were performed at 37 °C in a humidified incubator with 5% CO₂, unless otherwise specified. Before experiments, astrocytes were seeded in culture dishes and allowed to adhere for 12 h in complete culture medium consisting of DMEM supplemented with 10% fetal bovine serum and 1% penicillin-streptomycin. All cell handling procedures were conducted in a sterile laminar-flow biosafety cabinet that had been sterilized under UV irradiation for 30 min prior to use. Tyrode buffer solutions were sterilized by filtration through 0.22 μm membrane filters before application.

#### Supplementary Note S23: Preparation of Reagents

Fluo-4 AM working solution. Fluo-4 AM powder (50 μg per vial) was dissolved in 15.2 μL DMSO to prepare a 3 mM stock solution. The stock solution was aliquoted into 2 μL fractions, protected from light, and stored at −20 °C. For use, 2 μL stock solution was diluted in 1998 μL Ca^2+^-free Tyrode buffer to obtain a 3 μM working solution, which was freshly prepared before each experiment.

Thapsigargin solution (1 μM). A 10 mM thapsigargin stock solution in DMSO was diluted by adding 5 μL stock solution into 50 mL Ca^2+^-free Tyrode buffer, resulting in a final concentration of 1 μM. The solution was freshly prepared prior to use.

CaCl_2_-containing Tyrode buffer (4 mM). 22.2 mg CaCl_2_ was dissolved in 50 mL Ca^2+^-free Tyrode buffer to obtain a 4 mM CaCl_2_ solution. The solution was sterilized by filtration through a 0.22 μm membrane filter and stored at 4 °C for up to one week.

#### Supplementary Note S24: Establishment and validation of the active state (SOCE) of cells

Cell preconditioning. To remove extracellular Ca^2+^ and establish Ca^2+^-free conditions, the culture medium was first removed and cells were washed three times with 1 mL Ca^2+^-free Tyrode buffer, with 5 min incubation during each wash. Subsequently, cells were incubated with 1 mL Ca^2+^-free Tyrode buffer for 20 min to ensure complete removal of residual extracellular Ca^2+^.

Fluo-4 AM loading and de-esterification. After removal of the washing buffer, cells were incubated with 1 mL Fluo-4 AM working solution (3 μM) for 30 min in the dark to allow intracellular dye loading. The dye solution was then removed, and cells were washed three times with 1 mL Ca^2+^-free Tyrode buffer, each with 1 min incubation. To ensure complete intracellular de-esterification of the AM ester groups, cells were subsequently incubated in 1 mL Ca^2+^-free Tyrode buffer for 30 min in the dark.

Endoplasmic reticulum Ca^2+^ store depletion. For depletion of endoplasmic reticulum Ca^2+^ stores, imaging was initiated on the microscope stage. The buffer was replaced with 1 mL Ca^2+^-free Tyrode buffer containing 1 μM thapsigargin, and cells were incubated and continuously imaged for 10 min.

Induction of Ca^2+^ influx via SOCE. To trigger store-operated Ca^2+^ entry, 1 mL Tyrode buffer containing 4 mM CaCl_2_ was mixed with 1 mL of the previously collected Ca^2+^-free buffer from the dish to yield a final extracellular Ca^2+^ concentration of 2 mM. 1 mL of the mixed solution was then added back to the cells to induce Ca^2+^ influx through SOCE channels. Fluorescence imaging was performed for 10 min to monitor intracellular Ca^2+^ dynamics.

**Figure S21.**
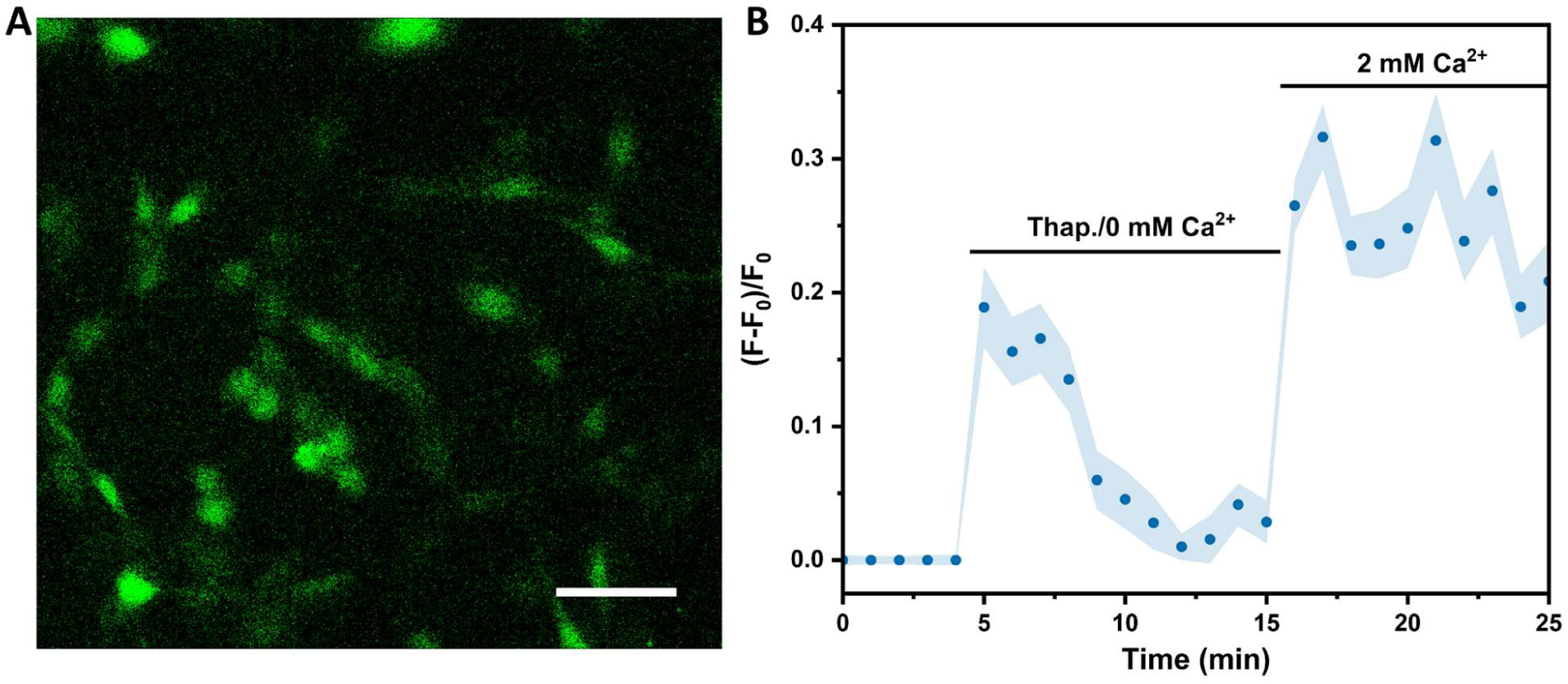
Validation of the active state (SOCE) of CTX TNA2 astrocytes. **(A)** Representative fluorescence image (scale bar: 50 µm) of CTX TNA2 astrocytes loaded with Fluo-4 AM. **(B)** Temporally continuous fluorescence imaging for monitoring intracellular Ca^2+^ dynamics of CTX TNA2 astrocytes under baseline control, endoplasmic reticulum Ca^2+^ store depletion (thapsigargin/0 mM Ca2+), and induction of Ca^2+^ influx via SOCE (2 mM Ca^2+^).

### Section 9: BISPIN microscopy for single-cell mapping of intrinsic stochastic magnetism

Cells (CTX TNA2 astrocytes, Hela, and NIH/3T3 fibroblasts) were cultured on plasmon-enhanced NV substrates. Imaging was performed under TIRF mode to maximize collection efficiency from near-interface NV emitters. Temporally continuous ODMR imaging was performed to capture stochastic magnetic fluctuations.

#### Supplementary Note S25: Digital ODMR mapping for multiple cell types

For BISPIN analysis, resonance splittings (Δ) were extracted from each NV emitter using the established ODMR fitting pipeline. Binary event classification was applied using the pre-established BISPIN decision threshold (Δ_th_), converting analog splitting values into positive and negative events. Positive-event fractions were calculated for each cell and imaging region to quantify magnetic activity. Baseline event distributions were first determined in the absence of cells to establish substrate-intrinsic noise levels. Paraformaldehyde-fixed cells were used as negative controls to assess contributions from passive cellular material. Live-cell measurements were then acquired under rest state and active state (SOCE).

In cases where cellular magnetic fluctuations are weak and lead to excessively low positive-event fractions, statistical significance can be improved by accumulating frames over time, effectively increasing the sampling of stochastic events. This frame accumulation approach ensures that the computed positive-event fractions more reliably reflect underlying cellular magnetic activity, allowing meaningful BISPIN analysis even under low-signal conditions.

**Figure S22.**
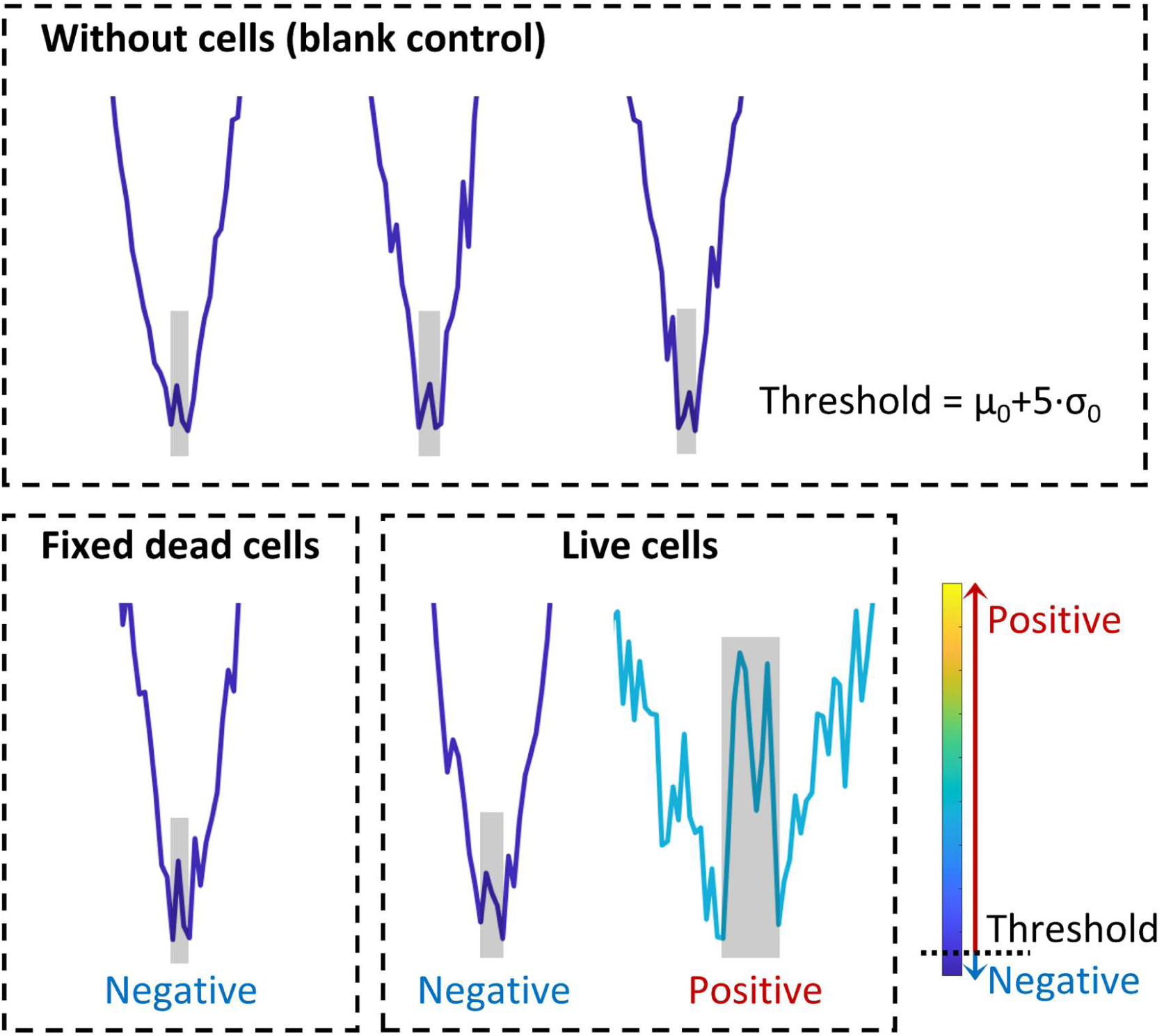
BISPIN thresholding for quantifying cellular stochastic magnetism. ODMR spectra are first acquired under blank control, defining the threshold Δ_t_*_ℎ_*= *μ*_0_ + 5 · *σ*_0_. For subsequent measurements on fixed (dead) and live cells, each ODMR-derived splitting value (Δ) is compared to the threshold Δ_th_. Events satisfying Δ ≥ Δ_th_ are classified as positive events, whereas events with Δ < Δ_th_ are classified as negative events.

**Figure S23.**
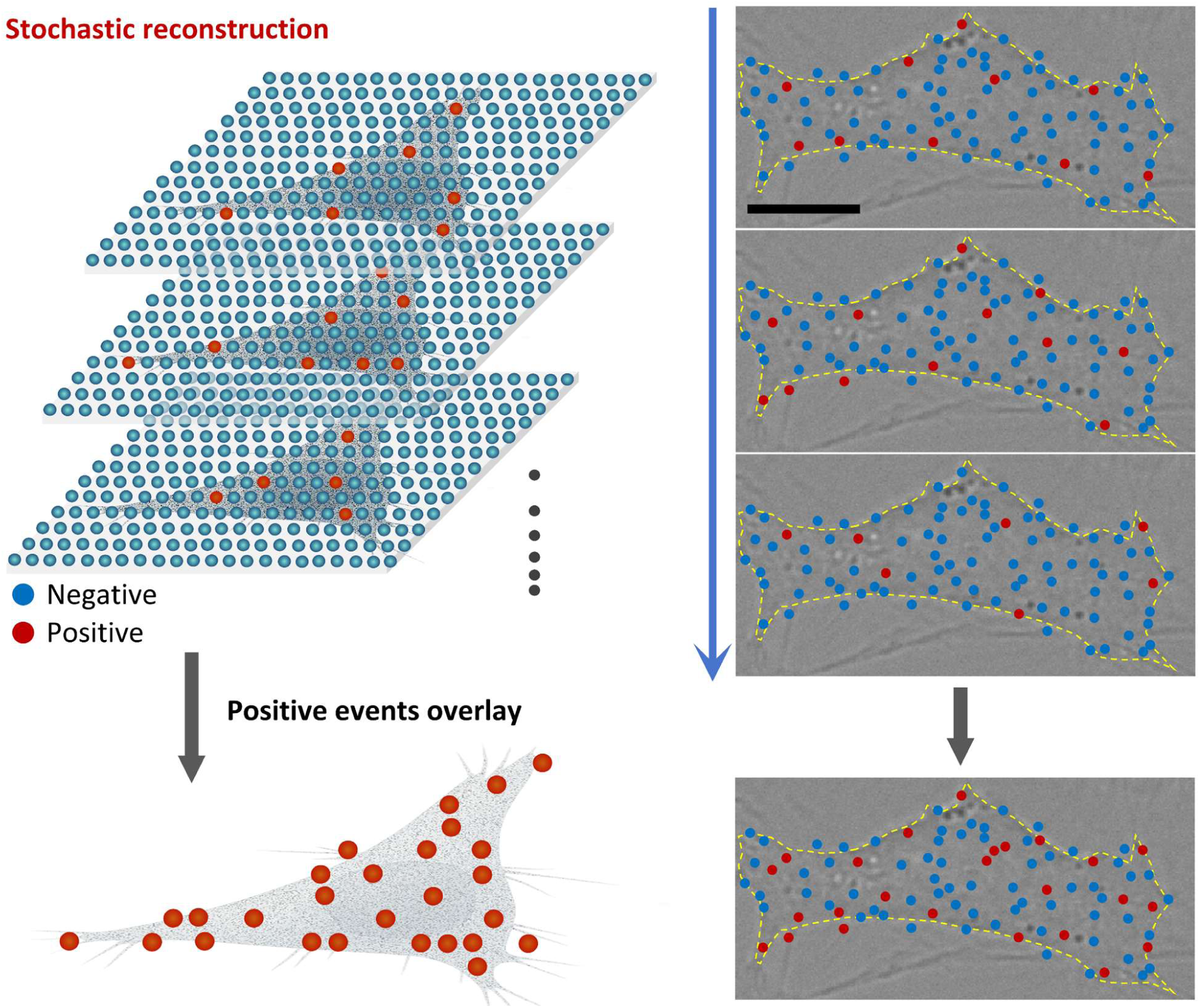
Cumulative BISPIN imaging for enhanced statistical significance. Cumulative BISPIN imaging of weak cellular magnetic fluctuations effectively increases the sampling of stochastic events, enabling the enhanced statistical significance and reliable BISPIN analysis under low-signal conditions.

#### Supplementary Note S26: Subcellular compartment-resolved BISPIN analysis

To analyze spatial heterogeneity, individual cells were segmented into whole-cell, peripheral edge, and interior cytoplasmic compartments based on optical images. Cell boundaries were delineated from bright-field images. Regions extending ±3 μm from the cell boundary were designated as the peripheral edge compartment, forming a 6 μm-wide area encompassing both sides of the cell outline. Regions located more than 3 μm inside the cell boundary were classified as the intracellular cytoplasmic compartment, whereas regions beyond 3 μm outside the cell boundary were assigned to the extracellular compartment. BISPIN positive-event fractions were computed for each compartment, enabling subcellular mapping of stochastic magnetic activity. For visualization, positive and negative events were color-coded and overlaid on corresponding optical images to generate spatially resolved event maps.

Event statistics, including positive-event fractions and spatial distribution patterns, were compared across cell types, functional states, and subcellular compartments. This workflow provides a label-free, contact-free, and statistically convergent method for single-cell mapping of intrinsic stochastic magnetic disturbances, with active cellular processes inferred from live-versus-fixed contrasts and SOCE-dependent modulation.

**Figure S24.**
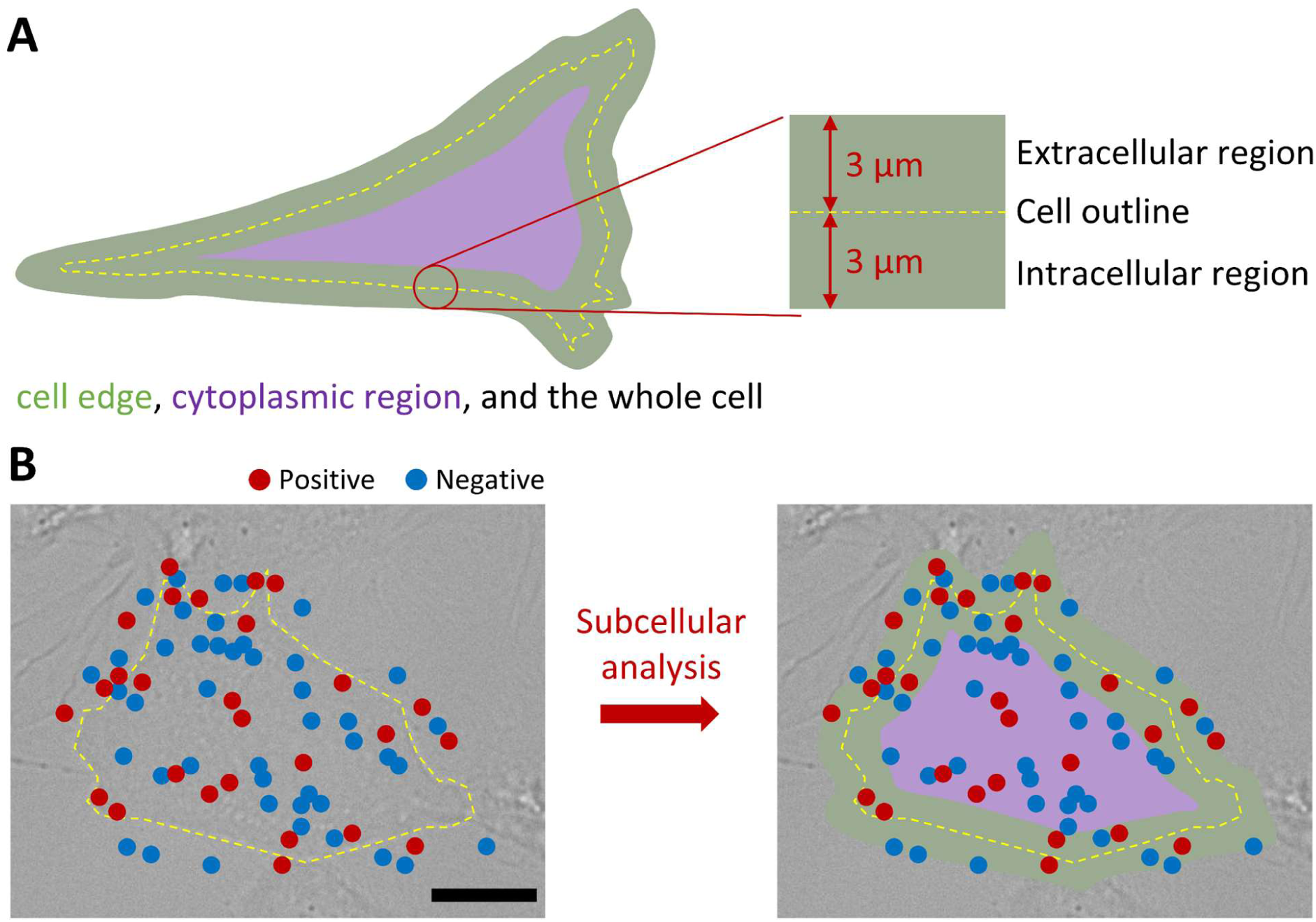
Subcellular compartment-resolved BISPIN analysis. **(A)** Schematic of subcellular compartment segmentation. Cell boundaries are defined from bright-field images (yellow dashed line). A ±3 μm region relative to the cell boundary is designated as the peripheral edge compartment, forming a 6 μm-wide band spanning both extracellular and intracellular sides. Regions located more than 3 μm inside the boundary were classified as intracellular cytoplasmic regions. **(B)** Subcellular analysis of BISPIN imaging enables quantitative calculation of compartment-resolved positive-event fractions and visualization of spatially heterogeneous magnetic activity within and around individual cells.

### Section 10: Machine-learning-assisted classification of single-cell bio-spin signatures

For each cell imaged by BISPIN microscopy, compartment-resolved positive-event fractions (whole-cell, peripheral edge, and cytoplasmic interior) were extracted as quantitative descriptors of stochastic magnetic activity. These values were used as feature vectors for supervised classification across six experimental groups: CTX TNA2 astrocytes, Hela cells, and NIH/3T3 fibroblasts under rest and active-state (SOCE) conditions.

Principal component analysis (PCA) was performed to visualize intrinsic structure in the multidimensional feature space. The first two principal components were used to assess clustering by cell type and functional state. Eight representative machine learning models were then implemented to evaluate classification performance: Logistic Regression, Support Vector Machine (SVM), Decision Tree, Random Forest, Gradient Boosting, AdaBoost, k-Nearest Neighbors (kNN), and a feedforward Neural Network. Models were trained on the compartment-resolved positive-event fractions and validated using standard cross-validation procedures. Receiver operating characteristic (ROC) curves were generated for each model, and the area under the curve (AUC) was calculated to quantify classification performance. Confusion matrices were used to assess misclassification rates and reproducibility across all six classes.

All machine learning analyses were conducted using Orange Data Mining (v3.34) with built-in preprocessing, feature scaling, and cross-validation workflows. This approach enabled quantitative evaluation of whether BISPIN-derived bio-spin signatures provide sufficient information to distinguish cellular identity and functional state without exogenous labels.

## Notes

### Competing Interest Statement

The authors have declared no competing interest.

